# Nuclease genes occupy boundaries of genetic exchange between bacteriophages

**DOI:** 10.1101/2023.03.23.533998

**Authors:** Zachary K Barth, Drew T Dunham, Kimberley D Seed

**Author notes:** These authors contributed equally to this work (Zachary K Barth, Drew T Dunham).

## Abstract

Homing endonuclease genes (HEGs) are ubiquitous selfish elements that generate targeted double-stranded DNA breaks, facilitating the recombination of the HEG DNA sequence into the break site and contributing to the evolutionary dynamics of HEG-encoding genomes. Bacteriophages (phages) are well-documented to carry HEGs, with the paramount characterization of HEGs being focused on those encoded by coliphage T4. Recently, it has been observed that the highly sampled vibriophage, ICP1, is similarly enriched with HEGs distinct from T4’s. Here, we examined the HEGs encoded by ICP1 and diverse phages, proposing HEG-driven mechanisms that contribute to phage evolution. Relative to ICP1 and T4, we found a variable distribution of HEGs across phages, with HEGs frequently encoded proximal to or within essential genes. We identified large regions (> 10kb) of high nucleotide identity flanked by HEGs, deemed HEG islands, which we hypothesize to be mobilized by the activity of flanking HEGs. Finally, we found examples of domain swapping between phage-encoded HEGs and genes encoded by other phages and phage satellites. We anticipate that HEGs have a larger impact on the evolutionary trajectory of phages than previously appreciated and that future work investigating the role of HEGs in phage evolution will continue to highlight these observations.

## INTRODUCTION

Genetic exchange, whether through sexual reproduction or horizontal transfer between organisms, is a major force underlying evolution. Typically, genes are selected for when they provide a benefit to their organism’s reproductive success. However, some genes and genetic elements select for their own propagation while being neutral or deleterious for the organisms that encode them. Gene drives are one such example of selfish genetic elements, and are defined as elements that bias the inheritance of certain genes beyond the 50:50 ratio predicted by Mendelian genetics (1). The capacity of gene drives to effect genetic change on a population level has made gene drives a topic of great interest, both for their technological potential and as one of the natural processes underlying evolution (2).

One of the most common forms of gene drive is homing endonuclease genes or HEGs. HEGs are ubiquitous, being found across all domains of life, and use an elegantly simple mechanism to drive their inheritance. HEG mobilization occurs first with sequence recognition by the DNA binding domain and subsequent cleavage of cognate loci that lack the HEG coding sequence by the endonuclease domain. Then, the HEG encoding sequence serves as a template for homology-directed repair, spreading the HEG coding sequence to its cognate gene locus (3–5).

Initially discovered within introns of fungal mitochondria, HEGs are now known to be highly abundant in bacteriophage genomes, where HEGs sometimes occur intronically, but are more frequently free standing genes (6). While Mendelian inheritance is not often associated with viral reproduction, coinfection of a single cell with distinct yet related phage isolates facilitates the recombination between phage genomes resulting in the production of hybrid progeny virions. HEGs have been shown to determine gene inheritance during phage crosses, ensuring that the HEGs themselves, as well as nearby sequences, are inherited by the hybrid progeny (7, 8). Beyond governing allele inheritance, HEGs can also serve as raw material for genetic innovation. There are notable examples of neofunctionalization among HEG-associated domains. For example, in *Saccharomyces cerevisiae*, the HO endonuclease responsible for mating-type switching shares a common ancestor with the WHO-element HEGs of *Torulaspora* and *Lachancea* yeasts (9). As another example, some isolates of the ICP1 group of phages possess the gene *odn*, which is HEG-derived and acts to protect the phage from subviral parasites (10).

HEGs have been extensively characterized in T4 and related phages (11–15), but interest in HEGs has not kept pace with the massive expansion of bacteriophage genomic research (16). T4-like phages have been noted as having a particularly high density of HEGs, and T4 itself stands out even among its relatives. 11% of T4’s coding sequence is occupied by HEGs (6). While the density of homing elements in T4-like phages is remarkable, it is not unique. ICP1, a more recently established but now extensively characterized (17–26) model phage that infects *Vibrio cholerae*, has been noted as having numerous putative HEGs (25). As ICP1 is frequently co-isolated with *V. cholerae* from cholera patient samples, several genetically distinct ICP1 isolates have been identified (19, 25, 27). These ICP1 isolates are highly similar but have been documented to encode variable genes. Given the high density of HEGs present in the ICP1 genome and the breadth of sequenced ICP1 isolates, ICP1 is an apt model organism for studying the impact of HEGs on phage genomics and evolution.

Here, we show that related HEGs are enriched in distinct phage lineages, including the ICP1-like phages, where HEGs are shown to associate with essential genes and allele exchanges similar to those described in T4-like phages. In ICP1, we show that this association between HEGs and essential genes often leads to the splicing of proximal genes, which we predict occurs in other phages as well. In many cases, HEGs bracket essential gene modules, forming what we have termed HEG-islands. HEG-islands show a striking level of nucleotide sequence conservation between phages that otherwise lack substantial nucleotide sequence similarity, suggesting that HEGs facilitate horizontal gene transfer between distantly related phages. We also observe sequence repetition within HEGs and domain shuffling between different HEGs, as well as between HEG and non-HEG genes. These architectural features highlight potential mechanisms for rapid HEG evolution, facilitating the emergence of novel genes within phages and phage-related mobile elements. Taken together, our observations show that HEGs are a driving force in bacteriophage evolution.

## MATERIAL AND METHODS

### Bacterial culture and strains

Experiments were conducted using *Vibrio cholerae* E7946 as a host (Table S1). Bacterial cultures were grown with aeration at 37°C in liquid LB medium. Cultures were supplemented with streptomycin (100 μg/mL) and ampicillin (100 μg/mL) for *Escherichia coli* cloning intermediates.

### Phage infection and isolates

All ICP1 isolates (Table S1) were stored as high titer stocks in STE buffer (100 mM NaCl, 10 mM Tris-Cl pH 8.0, and 1 mM EDTA). All bacteriophage infections were carried out at a multiplicity of infection of 2.5 when the host cell culture reached an OD_600_ of 0.3 and allowed to proceed until 16 minutes post-infection.

### RNA Isolation

*Vibrio cholerae* E7946 was grown and infected with ICP1 as described above. Infected samples were incubated at 37°C with aeration for 16 minutes, then mixed 1:1 with ice-cold methanol and vortexed. Samples were pelleted at 7,000 x g for 3 minutes at 4°C, washed with ice-cold 1x phosphate-buffered saline, and pelleted. The pellets were lysed in 200 μL of TRI Reagent (Millipore/Sigma) and incubated at room temperature for 5 minutes. After incubation, 40 μL of chloroform was added to each sample, samples were vortexed, and incubated at room temperature for 10 minutes. The TRI Reagent-chloroform mixture was then centrifuged at 12,000 x g for 10 minutes at 4°C to phase-separate the mixture. The aqueous phase was removed, added to a mixture of 110 μL of 2-propanol and 11 μL of sodium acetate pH 6.2, and mixed by vortexing. The precipitated RNA was pelleted from the mixture by centrifugation at 12,000 x g for 15 minutes at 4°C, washed twice with 75% ethanol, and the residual ethanol was evaporated from the washed pellets in a bead bath at 65°C. The RNA was resuspended in 20-30 μL of diethyl dicarbonate (DEPC) treated H_2_O and DNase treated according to the Turbo DNase kit (Thermo-Fisher). DNase-treated RNA concentration and purity were assessed by NanoDrop.

### Phage genomic DNA Isolation

100 μL of a high titer phage stock (>10^10^ plaque-forming units/mL) was treated with 1 μL of DNasel at 37°C for 20 minutes to degrade the un-encapsidated DNA and subsequently heat-inactivated at 65°C for 10 minutes. Phage genomic DNA was isolated following a modified version of the DNeasy gDNA Extraction Kit (Qiagen) as follows: 360 μL of buffer ATL and 40 μL of proteinase K were added to the DNase-treated phage, vortexed, and incubated at 65°C for 20 minutes. 8 μL of RNase A was added to the mixture, vortexed to mix, and incubated for 2 minutes at room temperature. 400 μL of AL and 400 μL of 100% ethanol were added to the mixture, the solution was vortexed, and the mixture was centrifuged through the spin collection column for 1 minute at 6,000 x g. 500 μL of wash buffer AW1 was added to the column and centrifuged at 6,000 x g for 1 minute. 500 μL of wash buffer AW2 was added to the column and centrifuged at 20,000 x g for 3 minutes to dry the column. The DNA was eluted from the column with 50 μL of dH_2_O after incubation at 65°C for 5 minutes.

### Splice junction validation

cDNA was synthesized using SuperScript III Reverse Transcriptase (Invitrogen) as per the manufacturer’s protocol. 1 μg of DNase-treated RNA was mixed with 100 ng of random hexamers and 1 μL of 10 mM dNTPs to a final volume of 13 μL. The mixture was heated at 65°C for 5 minutes and allowed to rest on ice for 1 minute. 4 μL 5x first strand synthesis buffer, 1 μL 0.1 M dithiothreitol, 1 μL of RNase inhibitor, and 1 μL of Superscript III reverse transcriptase were added to each reaction. Reverse transcription reactions were incubated at 25°C for 5 minutes, 50°C for 60 minutes, and 70°C for 5 minutes to inactivate the reaction. 1 μL of RNase H was added and the reaction was incubated at 37°C for 20 minutes, then incubated at 65°C for 10 minutes to heat-inactivate the enzyme. The inactivated reactions were purified with the Monarch PCR purification kit (NEB).

Standard Taq polymerase cDNA PCRs were carried out using 0.5 μL of cDNA as template DNA. Oligonucleotide primers were designed to amplify across the identified splice junctions from Magic-BLAST (28) analysis, and nested PCRs were carried out to amplify the desired products from the total cDNA synthesis products. Secondary PCRs were purified with the Monarch PCR and DNA Cleanup Kit (NEB), ligated into the pCR2.1-TOPO TA cloning vector (Invitrogen), and transformed into XL1 Blue *E. coli*. Transformants were screened by colony PCR and the splice junction sequences were determined by Sanger sequencing.

### MAGIC BLAST RNA-seq read mapping

Paired-end 150bp x 150bp RNA sequencing reads of *V. cholerae* infected with ICP1 16 minutes after infection were downloaded from the NCBI sequence read archive (PRJNA609114). Reads were trimmed to remove residual adapters and low-quality bases using Trimmomatic (v0.36) (29) and mapped to the ICP1 2006_Dha_EΔCRΔCas2_3 reference genome with MAGIC-BLAST (v1.5.0) using default settings. The resulting alignments were viewed with Integrated Genome Viewer (v2.3) (30) and regions of interest were manually identified for an intergenic drop in read coverage, corresponding to potential splice junctions for subsequent validation.

### Phylogenetic Analyses

All phylogenetic analyses comparing full phage genomes were carried out using ViPTree (v1.1.2) (31) with default settings. To generate phylogenies of proteins or domains, the sequences were first aligned using MAFFT (v7.505) (32) and the approximately-maximum-likelihood phylogenetic trees were generated using FastTree (v2.1.10) (33) with default settings. The resulting files were visualized and annotated with the iTOL (v6) (34) web browser.

### ICP1 and outgroup homing endonuclease domain prediction

Genomic sequences of each ICP1 isolate and the various outgroup phages were downloaded from NCBI using the Entrez e-utilities command line tools (for accession numbers, see Tables S1 and S2). Prediction of putative HEGs for all downloaded phages was initially performed with HMMER (v3.3.2) (35) using hmmsearch against the Pfam database with a maximum e-value cutoff of 0.001 to detect proteins with the Pfam domains indicated in Table S3. For proteins with T5orf172 domains, the MUG113 domain (PF13455) was also accepted, as the two profiles are highly related. ICP1 isolates were manually investigated using BLAST and EMBOSS-Needle to identify putative HEGs with frameshifts, deletions, or duplications in the coding sequence.

### Custom HMM profile generation

Representative amino acid sequences of domains from the ICP1 CapR and IPA-HNH domain-containing proteins were aligned with MAFFT and the resulting alignment was converted into a profile HMM with hmmbuild. The resulting HMM profile was searched against the UniProtKB database (Release 2021_02) with jackhmmer for 2 iterations, restricting results to bacterial viruses (taxid:28883). This search detected homologs outside of ICP1 phages, increasing the captured sequence diversity of the profile HMM. After two iterations, the resulting sequence alignment was downloaded as a profile HMM for further analysis.

### Gene neighborhood analysis

To detect phages that encode homologs to the ICP1 HEG T5orf172 and IPA-HNH families, we first generated a multiple sequence alignment of the representative HEGs of each family encoded by ICP1. For T5orf172 HEGs, this included representatives Gp53, Gp121, and Gp130. For HNH-IPA HEGs this included Gp59, the Gp114-115 fusion, and Gp209. The resulting alignment was converted to a PSSM, which was used as a query for two iterations of PSI-BLAST against the NCBI nr database, restricting the results to bacterial viruses (taxid:28883). This search resulted in a list of proteins with predicted similarity to the candidate HEG family. The genomes of phages encoding these proteins were downloaded from NCBI using Entrez e-utilities command line tools for further investigation. Any sequence less than 20 kb in length and all metagenome assemblies (most of the <20 kb length sequences), were removed from the dataset. Of the remaining sequences, the full genbank file with annotations of each phage was downloaded from NCBI (accessed August 2021). To standardize genome annotation across all of the phages, downloaded genomes were annotated with Prokka (v1.12) (36) using the Pfam-A v34.0 database for predicted functional annotations and novel coding sequence annotations predicted with Prokka were added to the genomes for downstream analyses.

These annotated genomes were then searched for proteins containing either T5orf172 or IPA-HNH HEGs with hmmsearch and gene neighborhoods were defined as the DNA sequence of 3,250 bp flanking the start and stop position of each T5orf172 or IPA-HNH HEG in the genome, extended to the bounds of any open reading frames which began or terminated beyond the allocated ± 3,250 bp sequence region. When this region exceeded the length of the genome, genomes were assumed to be circular and the search was expanded to the opposite end of the genome until the length criteria were met. The predicted protein domain content of the coding sequences encoded by the gene neighborhood was then annotated with hmmscan against the Pfam database, retaining predicted domains with a maximum e-value cutoff of 0.1. The predicted protein content of the gene neighborhoods was assessed based on these annotations, as indicated in Figure 2. Full gene neighborhoods analyzed in the study can be found in Tables S4 (T5orf172) and S5 (IPA-HNH).

### HEG Island and recombination prediction

The representative HEG islands reported in the manuscript were identified using BLASTn to detect large regions of high nucleotide similarity between less-similar phages. Putative HEG islands encoding phage structural genes identified in the gene neighborhood analysis were searched pairwise against distinct phages with BLASTn to identify representative examples of HEG island exchange. Regions of homology were marked on the gene graphs when they shared greater than 60 percent nucleotide identity over a length of at least 125 bp, with a maximum e-value cutoff of 1.0e^−5^. Representative identified HEG islands were graphed using DNA Features Viewer (37) and related sequences were shaded according to the BLASTn percent identity. HEG island functional annotation was performed with the HHpred web server against the Pfam database, allowing for functional prediction of phage genes that otherwise lacked annotation.

### DNA binding domain prediction and enumeration

To predict the DNA binding domains of the T5orf172 HEGs, we first identified the bounds of the predicted T5orf172 domain with HMMER. Because HEG nuclease domains are generally present on one terminus of the protein, we trimmed all amino acids from the T5orf172 domain to the nearest terminus of the protein, resulting in two sequences per protein, one predicted to encode a DNA binding domain and a second of the identified T5orf172 domain. These sequence lists were individually clustered with Cd-hit (v4.8.1) (38) to generate clusters of proteins at 90% sequence identity to remove highly related sequences and the clusters were converted into a multiple sequence alignment with MAFFT. Approximately maximum-likelihood phylogenetic trees were generated from the multiple sequence alignments with FastTree.

To predict the DNA binding domain(s) of the truncated sequence predicted to contain the DNA binding domains, we first used hmmsearch to identify proteins containing either the CapR domain (custom profile) or DUFs723 and 4397 (PF05265 and PF14344, respectively), both of which are predicted to encode Zn-finger DNA binding domains similar to the CapR domain. Sequences that were predicted to contain a CapR domain with a maximum e-value of 1.0e^−0^ were annotated as CapR DNA binding domains. Sequences which did not meet this threshold but were predicted to encode either DUF723 or DUF4397 were annotated as accordingly. Sequences that matched none of the three domains were further investigated with HH-suite3 (v3.2.0) (39), first aligning sequences against the UniRef30 (2022_02) target database with two iterations to generate a representative a3m alignment. This alignment was then searched against the Pfam database using HHsearch. DUF723 or DUF4397 domains identified from HHsearch were annotated accordingly, but were the number of domains present was not quantified given the difference in sensitivity between HHsearch and hmmsearch. Manual investigation of the HHsearch results also revealed the presence of RepA_N domains present in some T5orf172 proteins, leading to the identification of the putative *E. coli* satellites.

### Prediction of putative E. coli phage satellites

The DNA sequence of each of the three phage-encoded RepA_N-T5orf172 proteins was searched using BLASTn against the NCBI nt database to identify proteins with high primary sequence similarity, indicative of potential recombination events between phage and bacteria. Genomes encoding these homologous sequences were downloaded and the surrounding 12,500 bp on either end of the homologous sequence was extracted. The three putative satellite sequences that contained RepA_N domain proteins with the highest levels of homology to the previously identified phage RepA_N-T5orf172 protein were manually investigated for features indicative of satellite function, using a combination of BLAST and the HHpred webserver for functional prediction.

First, the three sequences were searched against one another using BLASTn, identifying large regions of gapped homology, approximately 13 kb in length. The bounds of these regions were manually investigated for inverted repeat sequences, indicative of integrase-bound attachment sites. Genes encoded between the attachment sites were then functionally annotated using the HHpred web server against the Pfam database to identify the prediction function of the encoded proteins. This search revealed two types of integrases and two distinct attachment sites present on the three representative sequences. Using these identified sequences, we then subjected the remaining 393 putative satellite-encoding contigs to a series of BLASTn searches to determine whether the observed integrases and attachment sites were present. Any putative element that contained less than 10 kb of noncontiguous homology to one of the three representative elements was discarded, leaving 224 potential satellites. These remaining satellites are summarized in Table S6, where we predict the putative attachment sites, which family of integrase is encoded by the satellite, the bounds of the satellite, and the sequence of the attachment sites. These findings are not meant to be comprehensive but provide a list of potential sequences for further investigation.

## RESULTS

### ICP1 phages encode diverse families of HEGs

Previously, numerous putative HEGs were identified in ICP1 phages (25). While a few of these genes contained domains previously linked to HEG activity, such as HNH-3 and LAGLIDADG, approximately half of these putative HEGs encoded the less-characterized T5orf172 domain (Figure 1A). T5orf172 is a subfamily of GIY-YIG endonucleases, bearing a similar core motif, and in at least one case, T5orf172 nuclease activity was shown to be dependent on a glutamate residue whose catalytic function was inferred from multiple sequence alignments with biochemically characterized GIY-YIG endonucleases (10). The broader GIY-YIG family has been associated with homing activity in phages, and the majority of HEGs in T4 possess a GIY-YIG domain (Figure 1A) (6, 14). To determine if a similar enrichment of T5orf172 HEGs occurs across the ICP1 isolates and phylogenetically related ICP1-like phages, we bioinformatically identified the putative HEGs across this group of phages, using profile HMM predictions to identify genes that encode putative HEG domains according to the Pfam database. This analysis confirmed an abundance of T5orf172 HEGs in ICP1 phages and demonstrated the diversity of HEGs present in otherwise phylogenetically related phages. Beyond having HEG-associated domains, ICP1’s putative HEGs exhibit several genomic signatures characteristic of HEGs: their presence is variable between ICP1 isolates (Figure 1A) (11, 12); when present, the genes are unstable due to frameshifts, variable stops, or in-frame deletions (11) (Figure 1A); they often occur near or inside essential phage genes (40); and in the case of ICP1’s T5orf172 genes, there is a high level of nucleotide similarity between related genes, suggesting the amplification and diversification of a common ancestor throughout ICP1 genomes over time. While the homing activity of ICP1’s putative HEGs has not been experimentally verified, these genomic signatures provide strong evidence for homing. For the sake of concision, we will refer to these putative HEGs simply as HEGs.

**Figure 1.**
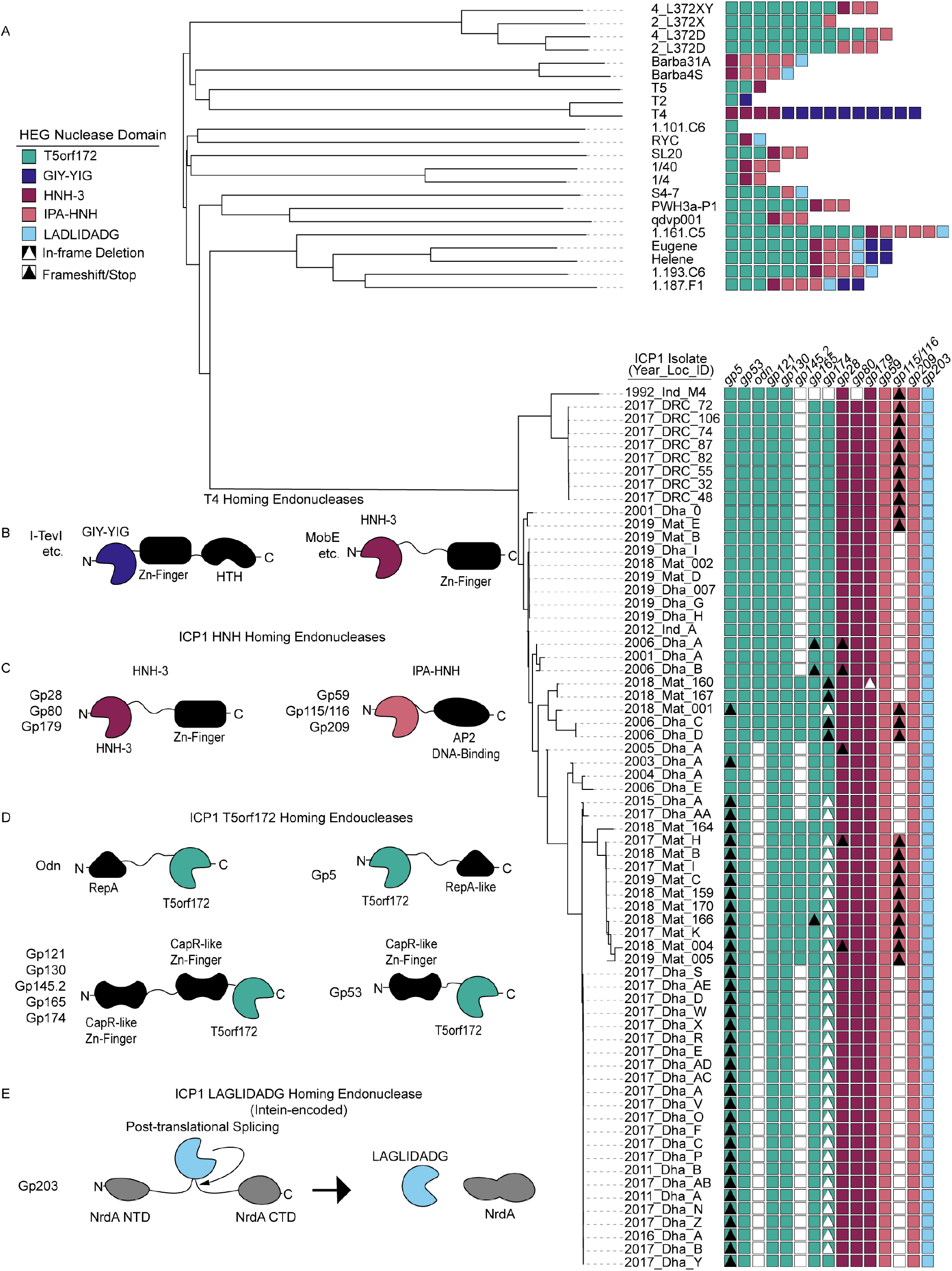
ICP1 phages encode many homing endonucleases and are enriched for T5orf172 genes. (A) A phylogeny of the 67 ICP1 isolates (bottom clade) and various outgroup phages (top clade). ICP1 isolates are named according to the year of isolation, country of isolation, and an identifying character to differentiate phages co-isolated in the same year and location (e.g., 2017_Dha_A = 2017 isolate from Dhaka, Bangladesh, isolate A). A VipTree alignment of all genomes of interest and subsequent HMM profile predictions of homing endonuclease genes in each representative genome shows enrichment of T5orf172 homing endonucleases in the clade of ICP1 isolates compared to most outgroup phages. Notably, highly related ICP1 isolates encode variable homing endonuclease genes, with frequent frameshifting (black triangle), in-frame deletion (white triangle), and lack of HEG altogether (white box) represented. These predictions are likely underestimates of the full HEG repertoire among these phages due to the genetic drift of HEGs impeding functional predictions. (B-E) The variable homing endonuclease domain architecture between T4-encoded (B) and ICP1-encoded homing endonucleases (C-E). The predicted DNA binding domains are shown as black shapes and the predicted nuclease domains are colored according to the legend in (A). Apart from the LAGLIDADG HEG *gp203*, which is intein encoded and post-translationally spliced out of the functional NrdA protein, each of the represented domain architectures shares the general pattern of one terminus encoding at least a single DNA-binding domain and the opposite terminus encoding the nuclease effector domain.

ICP1 phages also were found to encode HEGs belonging to numerous other HEG families (Figures 1B, 1C, 1D, and 1E). ICP1 isolates encode some HNH-family endonucleases with similar domain architecture to T4 HNH HEGs (Figures 1B and 1C). Within ICP1, *gp28, gp80*, and *gp179* each encode an N-terminal HNH-3 domain and a C-terminal zinc-finger domain similar to the Mob class of HEGs from T4 (41). Additionally, we identified three ICP1 genes, *gp59, gp115*, and *gp209*, with domains related to, but distinct from, the typical HEG-associated HNH-3 domains. While these domains are too divergent from canonical HNH-3 domains for profile predictions using the HNH-3 Pfam profile, HMM-HMM searches using HHSearch identified putative N-terminal HNH domains in each of the three proteins, and both *gp59* and *gp209* encode C-terminal AP2 DNA binding domains. Although *gp115* does not encode any predicted DNA binding domain, *gp116* immediately downstream encodes an AP2 domain. We reasoned that these two genes likely represent an inactivated HEG due to a frameshift, as the fused *gp115-gp116* contains the expected domain architecture, and we performed all subsequent analyses with this fused protein sequence. In support of these genes as atypical HNH HEGs, this combination of an N-terminal HNH domain and a C-terminal AP2 domain has been previously identified as a family of HEGs in protists, cyanobacteria, and phages (42). Based on the domain architecture of these genes and the previous characterization of related genes, we conclude that they represent a class of HEGs that exists within the larger HNH superfamily and will refer to them as **<u>I</u>**CP1-like **<u>P</u>**hage **<u>A</u>**P2 domain associated HNH-domain HEGs or IPA-HNH HEGs.

### T5orf172 and IPA-HNH HEGs are enriched in diverse phage lineages

As phage HEG research has been largely dominated by the study of T4-like phages, and T4 does not possess T5orf172 genes or IPA-HNH genes, we searched broadly among phages for HEGs homologous to those in ICP1, hoping to broaden our understanding of HEG diversity within and across phage genomes. Besides their abundance in the ICP1 isolates and closely related phages, both T5orf172 and IPA-HNH genes are enriched in multiple taxonomically diverse lineages. T5orf172 genes are abundant in the Plateaulakeviruses, a clade of *Aeromonas sp*. infecting phages, T5-like phages, and the dsDNA tail-less phages of *Autolykiviridae* (Figure 2A). IPA-HNH coding genes are abundant in separate lineages, namely the marine Barbaviruses, and the *Pseudomonas sp*. infecting Flaumdraviruses (Figure 2B). The reasons for the enrichment of specific phage taxa with specific HEG families are unclear. Possible explanations include specific biological traits, such as a phage’s ability to exclude co-infecting phages, and early invasion into a phage lineage by a specific HEG class.

**Figure 2.**
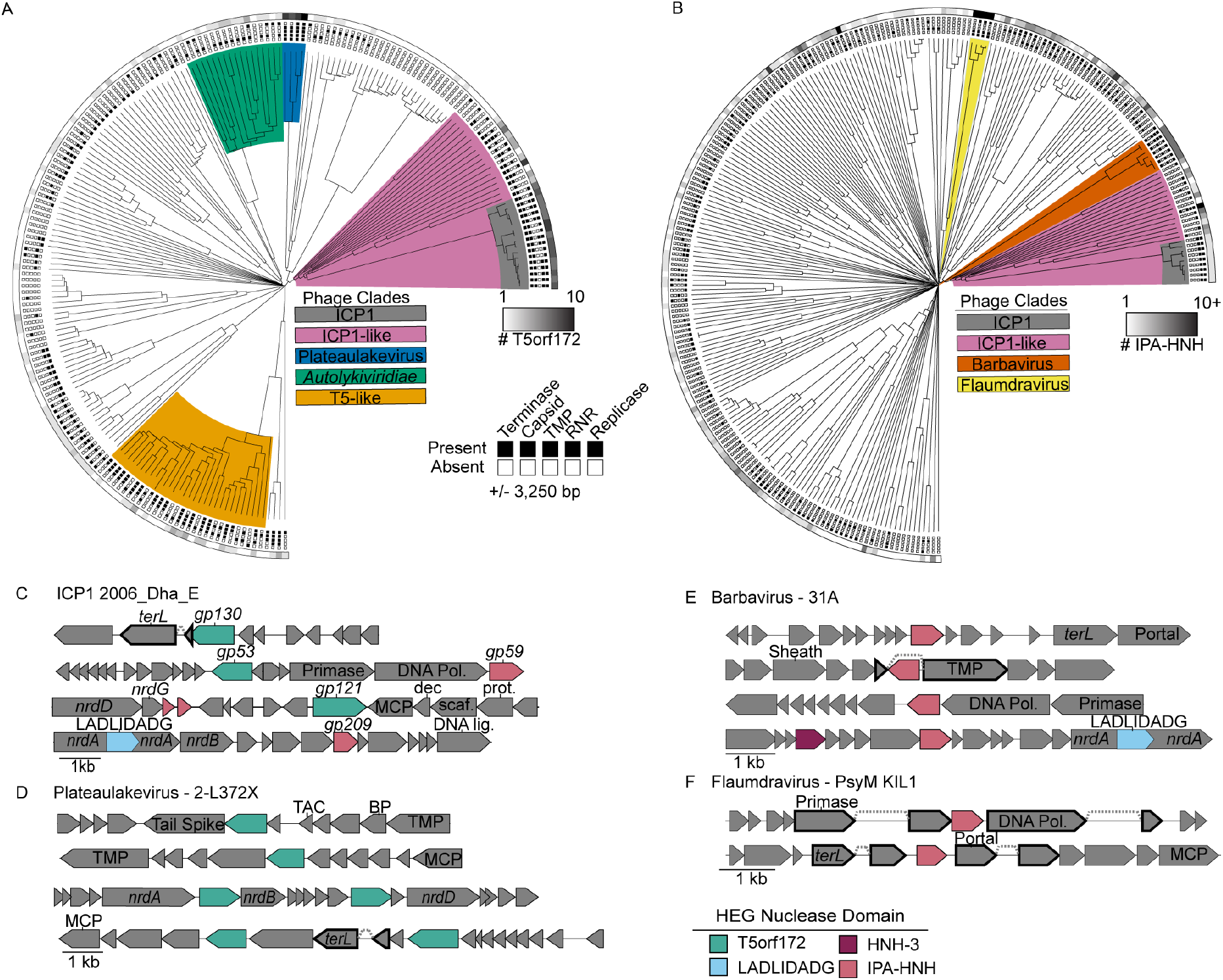
HEGs cluster near essential genes in diverse phage lineages. (A, B) T5orf172 (A) and IPA-HNH (B) HEGs are often localized in gene neighborhoods that include essential phage genes. Candidate phage genomes containing homologs to ICP1 representative nucleases were identified by PSI-BLAST and the gene neighborhoods containing the ± 3,250 bp flanking each predicted HEG present were annotated for predicted function. Gene neighborhoods including genes predicted to encode a phage terminase, capsid, tape measure protein (TMP), ribonucleotide reductase (RNR), or replicase component were indicated with a black box at the end of the leaf. The number of putative HEGs in each genome is represented by the grayscale bar surrounding the tree, with all phages encoding at least one representative HEG. Clades with a high density of HEGs discussed throughout the manuscript are highlighted accordingly. (C-F) Representative gene neighborhoods for T5orf172 (C, D) and IPA-HNH (C, E, F) HEGs show the co-localization of HEGs with essential phage genes. Validated and predicted exon splicing junctions are indicated by a dashed line connecting two genes, which are outlined in bold. Genes encoded by ICP1 are shown with their gp number, as referenced in Figure 1. Colored boxes next to phage names indicate the phage clade as colored in (A) or (B). Functional predictions for phage genes are based on the Pfam domain predictions using hmmsearch. Domain key: *terL* = large terminase, DNA Pol = DNA polymerase, *nrdA/B/D* = ribonucleoside-triphosphate reductase, MCP = major coat subunit, Dec = decoration protein, Scaf = scaffold, Prot = scaffold protease, TAC = tail assembly chaperone, BP = baseplate, TMP = tape measure protein.

We next constructed a phylogenetic tree of T5orf172 domain sequences to examine if the relationship between T5orf172 proteins also suggested expansion within specific phage lineages (Figure S1). Generally, T5orf172 sequences are more related to sequences from the same phage lineage, but we also found evidence for some inter-lineage exchange of T5orf172 genes (Figure S1). A small number of T5orf172 domains from ICP1 and ICP1-like phages occur within a clade comprised of sequences from T5-like phages, while about half of the T5orf172 genes in ICP1 form a distinct clade the other half are more dispersed. These results indicate that HEG expansion within specific phage lineages is primarily responsible for HEG distribution, but there is an appreciable level of horizontal HEG transfer between different phage lineages.

### HEGs cluster near essential genes in diverse phage lineages

Previous work investigating HEGs in phages has shown that HEGs often integrate near or within essential genes or operons (6, 40, 43–45), including those encoding structural genes, the terminase, ribonucleotide reductases, and the replisome. Using the previously identified genomes that encode T5orf172 and IPA-HNH HEGs, we performed a gene neighborhood analysis surveying the genes encoded within the flanking 3,250 bp of the detected HEGs, aiming to assess whether a similar association occurs between the HEG families in our dataset and their integration sites (Figure 2A and 2B). Consistent with previous work investigating HEGs in phages, we find that both T5orf172 and IPA-HNH coding genes often occur near or within essential genes and operons, including those for structural genes, terminase, ribonucleotide reductase, and the replisome (Figure 2C-F). Of the 468 T5orf172 HEG gene neighborhoods analyzed across 204 phage genomes, 35% of the neighborhoods contain a predicted terminase gene, 33% encode predicted ribonucleotide reductase genes, and 31% encode predicted capsid genes (Figure 2A, Table S4). We see a similar density but weaker overall trend in the association of IPA-HNH HEGs (n = 431 HEGs across 240 genomes) with these essential loci. The 431 IPA-HNH HEGs are most frequently found in gene neighborhoods predicted to encode DNA polymerase (29%), ribonucleotide reductase (18%), and terminase genes (14%). Despite this decreased relative frequency of IPA-HNH association with the indicated marker genes, specific clades of phages, such as the Barbaviruses, show a strong association between the HEGs and the indicated gene families (Figure 2B, Table S5). Notably, not all phages identified in this analysis are expected to encode all of the indicated genes due to differences in the phage morphology and lifestyle. One such representative example, the tail-less *Autolykviridae* would be expected to encode neither a tape measure protein nor a terminase, and many phages rely on cell-encoded replisomes rather than encoding their own. In cases such as these, there may be selection for HEG integration in different loci essential for a given phage lifestyle. The absence of these features in some genomes results in an underrepresentation of the frequency of HEG integration proximal to the indicated genes.

When these gene neighborhoods were examined more closely, many interesting trends were observed. We often found HEGs of other classes near T5orf172 or IPA-HNH HEGs (Figure 2C and 2E), suggesting these loci may be favorable for HEG acquisition or retention. While the majority of HEGs observed appear to be free-standing genes, we did observe numerous cases where other genes were interrupted by a HEG sequence. At least 8 of the 20 LAGLIDADG endonucleases within our dataset were predicted to be intein-encoded within a *nrdA* gene, similar to that of ICP1 (Figure 2E). Another frequently interrupted gene encodes the tape measure protein (TMP), which serves as a scaffold for tail tube assembly (46). ICP1’s own TMP is interrupted by a HEG, consistent with other documented examples of intron-associated HEGs and spliced transcripts (47–49).

To determine the splicing of ICP1’s interrupted TMP gene forms a full-length TMP gene, we analyzed previous RNA sequencing data generated during ICP1 infection (24, 50) with the NCBI Magic-BLAST tool, which is designed for detecting candidate intronic sequences from RNA sequencing data sets (28). From this analysis, we detected the splicing of the multiple ICP1 transcripts (Figure 3). To confirm the splicing of the ICP1 TMP and determine the full exonic sequence, we used reverse transcriptase PCR (RT-PCR) of RNA isolated during phage infection coupled with Sanger sequencing. The ICP1 TMP coding transcript is comprised of four exons and three introns, with the middle intron typically encoding an HNH-3 HEG (Figure 3A). Remarkably, the oldest ICP1 isolate sequenced thus far, ICP1 1992_Ind_M4 (ICP1 M4), encodes variable intronic sequences while maintaining the splice junctions and undergoing only minimal changes to the surrounding exonic nucleotide sequence (Figure 3B). Instead of a single HNH-3 HEG occupying the middle intron, ICP1 M4 has two HNH-3 HEGs, one in the first and third intron, while the middle intron is empty (Figure 3A). Each of the HEGs shares approximately 30% amino acid identity with another, suggesting this locus tolerates the integration of distinct HEGs. HEG-driven gene conversion is well documented, but this example from ICP1 shows that the HEG content of introns can vary while the surrounding sequence context is nearly unchanged, adding another evolutionary route for the diversification of homing loci.

**Figure 3.**
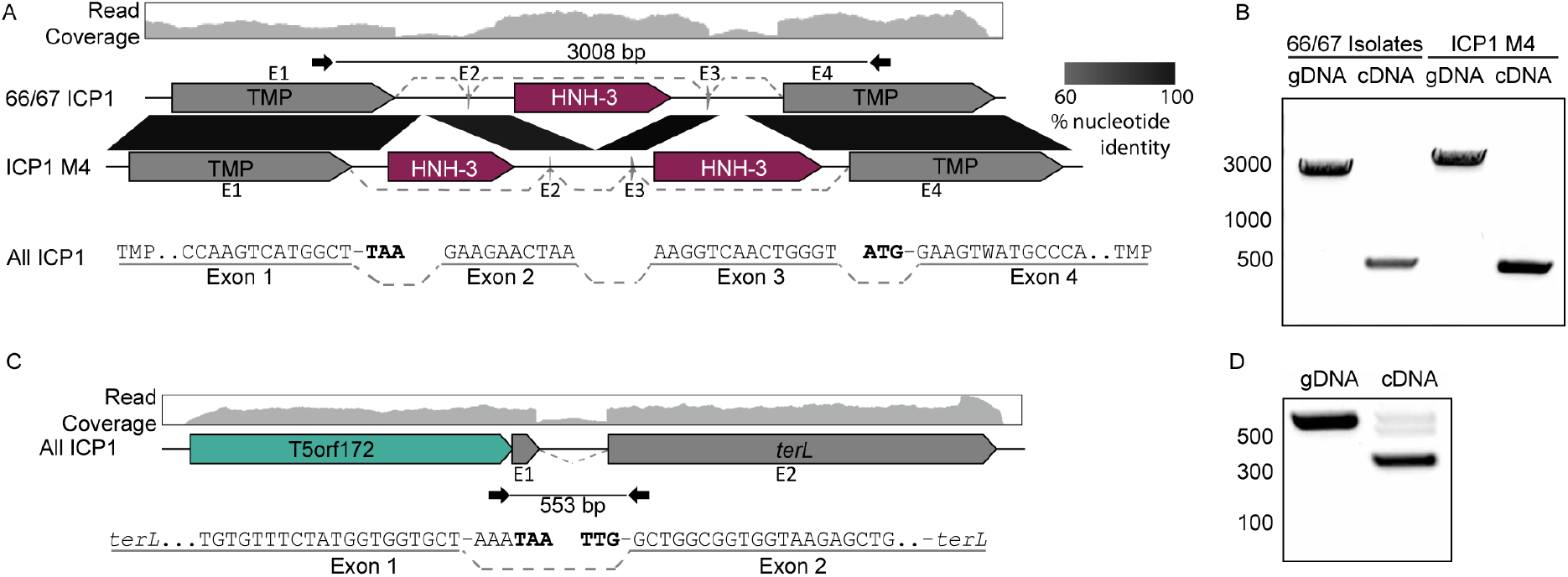
ICP1 isolates use conserved transcript splicing to express fragmented essential genes. (A, C) Gene graphs representing the splicing architecture of the ICP1 tape measure (A) and large terminase (C) as detected by RNA sequencing and RT-PCR. 66 of the 67 ICP1 isolates share a similar architecture, with a single HNH-3 HEG between the two TMP fragments (Top gene graph). A single ICP1 isolate, ICP1_1992_M4 (ICP1 M4), encodes two intronic HNH-3 HEGs in the same locus, both unique from the HNH-3 HEG found in the majority of ICP1 isolates (bottom gene graph). Despite the considerable difference in intronic sequence, all ICP1 isolates undergo differential splicing to excise unique homing endonuclease genes from TMP transcripts (A), resulting in nearly identical TMP sequences. Percent nucleotide identity between the conserved sequences is indicated by the black parallelogram between the gene graphs. All ICP1 isolates share conserved splicing of the large terminase (*terL*) (C), which is neighbored by a T5orf172 HEG. RNA-seq read mapping of total RNA isolated from ICP1 2006_Dha_E (top read coverage graphs aligned to a gene graph of the TMP (A) or terminase (C) coding region) shows a distinct drop in reads in intronic regions compared to that of the exonic regions. The splice junctions of the exonic transcript are represented below the gene graph and exons are labeled E1-4 (TMP) or E1-2 (*terL*), with dashed lines representing sequences joined by splicing. Nucleotides highlighted in bold are the predicted start and stop codons of the annotated coding sequences on the gene graphs, which are removed during the splicing process. (B, D) Agarose gels of PCR products generated following PCR across the splice sites of the TMP (B) or *terL* (D) sequence from gDNA and cDNA confirms splicing present in cDNA samples. Selected size markers from 2-Log DNA Ladder are indicated by the numbers to the left of the gels in base pairs, Sanger sequencing of cDNA products confirmed the exon mapping represented in (A) and (C). Black inward-facing arrows represent the primers used for both the cDNA and gDNA PCR reactions and the indicated size of the gDNA product is indicated in base pairs.

The Magic-BLAST analysis of RNA-seq data also revealed the splicing of ICP1’s terminase gene (Figure 3C), and the splicing of the terminase transcript was confirmed through RT-PCR analysis (Figure 3D). While phage functional annotation pipelines identified the larger C-terminal exon as encoding a full-length large terminase, the N-terminal exon had not been given a predicted function, as is commonly the case for short phage genes. Notably, ICP1 encodes a T5orf172 HEG adjacent to the terminase N-terminal exon rather than inside of the terminase intron (Figure 3C). Such an arrangement between introns and HEGs has been observed previously in T3 and other T7-like phages, as well as a cyanophage, and is described as collaborative homing (51, 52). In cases of collaborative homing, HEG endonucleolytic cleavage occurs at the intron insertion sequence, promoting the acquisition of both the HEG and the intron together. In such a scenario, it is not clear how the arrangement for collaborative homing would arise in the first place. An alternative hypothesis is that these collaborative homing loci occur due to HEG turnover, leading to the acquisition of a new free-standing HEG, and the purging of the intronic HEG, while leaving the intron splice junctions intact.

### HEGs bracket essential genes and operons forming ‘HEG-islands’

Often, multiple HEGs are encoded near the same essential locus. This suggests that similar to collaborative homing between free-standing HEGs and introns, collaborative homing may also occur between neighboring HEGs with different specificities. Cooperation between HEGs could increase the mobility of each HEG individually while also helping to avoid replacement by foreign homing loci. We observed many instances of essential genes, and in some cases, entire core operons, flanked on both ends by HEGs (Figure 4, Figure S2). This arrangement is suggestive of a mechanism wherein the flanking HEGs serve as vehicles to mobilize the essential loci, which are cargo of a complex homing region. In line with this model, we have named these regions HEG-islands. HEG-islands show remarkable conservation between phages on the nucleotide level (Figure 4, Figure S2). In phages that lack extensive nucleotide similarity across most of their genomes, we find HEG-islands with nucleotide identity greater than 90%, an incredible level of conservation for cargo mobilized through a recombinational repair homing mechanism. Many HEG-islands encode small genes of unknown function. Some of these small genes appear to be fragments of essential cargo with nucleotide homology to full-length genes (Figure 4B and 4C). These remnants may continue to serve a function for the HEG-island by acting as recombination substrates for future homing events, and recombination with intact essential genes may allow these fragments to serve as a reservoir of gene diversity.

**Figure 4.**
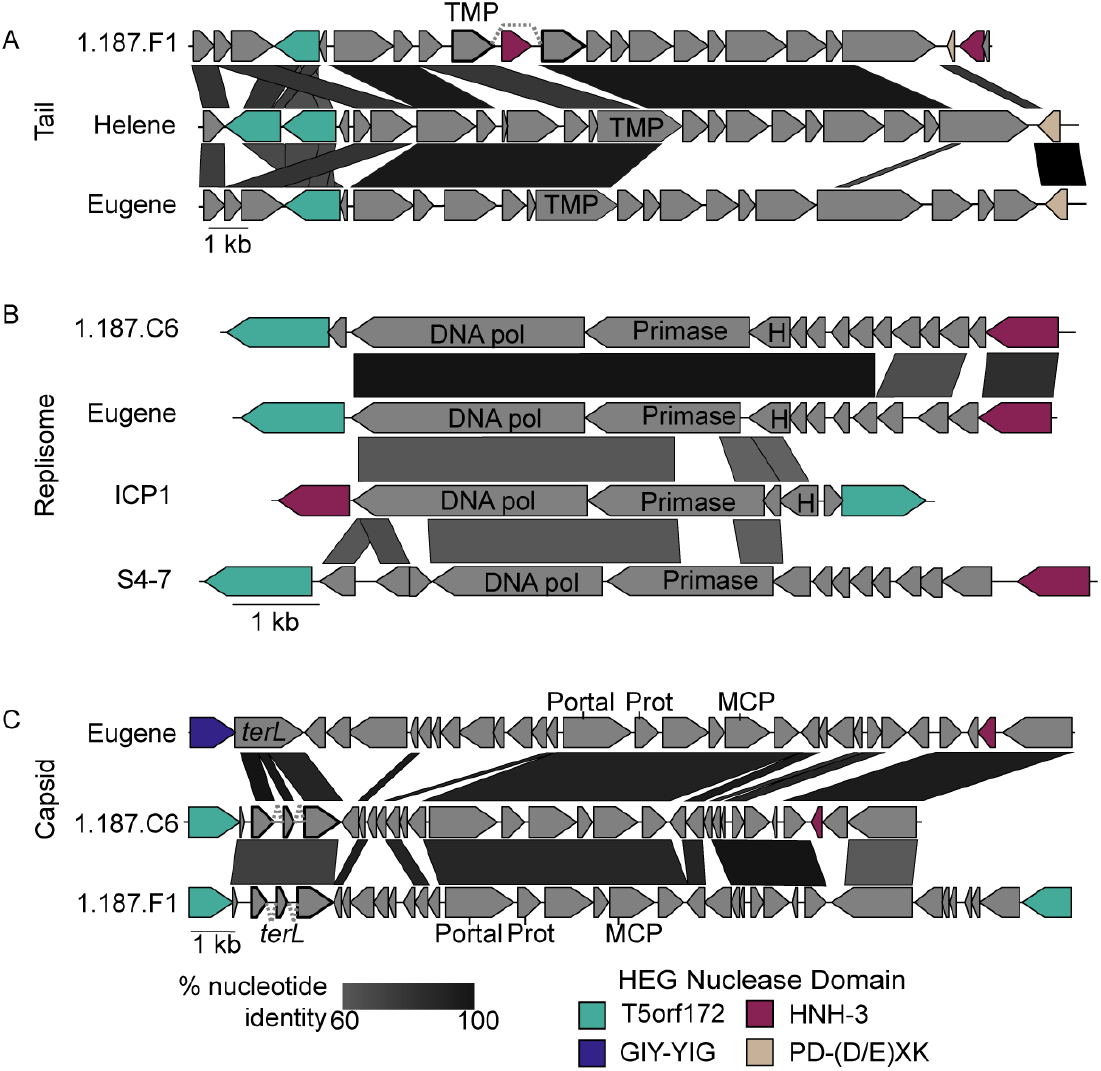
HEG-islands bracket regions of essential gene cargo in phages. (A-C) Gene neighborhoods are shown that contain HEGs in proximity to genes involved in phage tail morphogenesis (A), phage genome replication (B), or phage capsid morphogenesis (C). Examples of greater than 60% nucleotide identity between the phages are indicated by black-shaded parallelograms between the gene graphs. Gene fragments predicted to splice are outlined in bold, with gray dashed lines connecting the exons. Genes are colored according to the HEG nuclease domain and labeled according to the predicted gene function, as determined by HHsearch domain prediction. Domain key: *terL* = large terminase, DNA Pol = DNA polymerase, *nrdA/B/D* = ribonucleoside-triphosphate reductase, MCP = major coat subunit, Prot = scaffold protease, TMP = tape measure protein, H = RNase H.

The HEG content of related HEG-islands is unstable. While the related HEG-islands we observed typically had HEGs in the same locations, these cognate HEGs could be homologs with shared nucleotide identity, HEGs of the same class without obvious nucleotide similarity, or HEGs of entirely different classes (Figure 4, Figure S2). In one representative example, the tail operons (Figure 4A), we observed the replacement of the commonly occurring HNH-3 HEG with a PD-(D/E)XK HEG, a family HEGs found in bacteria (53, 54) but not found in ICP1 or most ICP1-like phages. HEG interchangeability may come about because multiple HEGs have evolved symbiotic relationships with related cargo genes, increasing the inheritance of the HEGs while concurrently affecting the fitness of the host through maintenance or turnover of essential gene modules. Given the essential functions of their cargo and their presence in widely diverged phages, HEG-islands appear to be a major force of phage evolution.

### HEGs and HEG-related proteins display repetitive domain architecture and domain shuffling

We next wanted to assess the domain architecture of the T5orf172 HEGs to gain more insight into their evolution. Previously, it was recognized that the DNA binding domains of ICP1 T5orf172 HEGs have homology to CapR (55), a transcriptional repressor of ICP1’s capsid expression encoded by phage-inducible chromosomal island-like elements (PLEs). PLEs are sub-viral parasites that exploit the ICP1 life cycle for their own mobilization (21–24, 26, 50, 55–58), inhibiting ICP1 progeny formation. This antagonism is frequently met with counter-adaptation by the phage (10, 18, 20, 59), providing a model system for the study of subcellular molecular co-evolution. The CapR DNA-binding domain was shown to be a Zn-finger domain with a CxxC…CxC motif ((24). Using a custom HMM profile representative of the CapR DNA binding domain from all known PLEs, we found that many of the T5orf172 genes in our dataset are predicted to encode at least one CapR domain (95/229 T5orf172 HEGs, 41.5%), providing further support to the association between the CapR and T5orf172 domains. Of those that did not encode a predicted CapR domain, one of two other highly related DNA binding domains, DUF723 or DUF4397, was almost always present. All three profile HMMs are distinct from one another, but each contains a similar Zn-finger domain and we anticipate that these domains each provide DNA binding activity to the HEG. In total, the vast majority of the identified T5orf172 HEGs (200/229, 87.3%) contained at least one of the three Zn-finger domains (Figure 5). Interestingly, we found that many T5orf172 encoding genes encode multiple distinct Zn-finger domains (Figure 5B). This repetitive domain organization is characteristic of class V repeat proteins. Class V repeat proteins can be described as ‘beads on a string’ as they consist of repetitive, independently folded and functional repeat domains connected by linker sequence (60). Target recognition is then an emergent property of the different domains acting together to provide specificity of DNA binding. These repeat domains may act cooperatively to provide such specificity to T5orf172 HEG DNA binding or to regulate cleavage activity.

**Figure 5.**
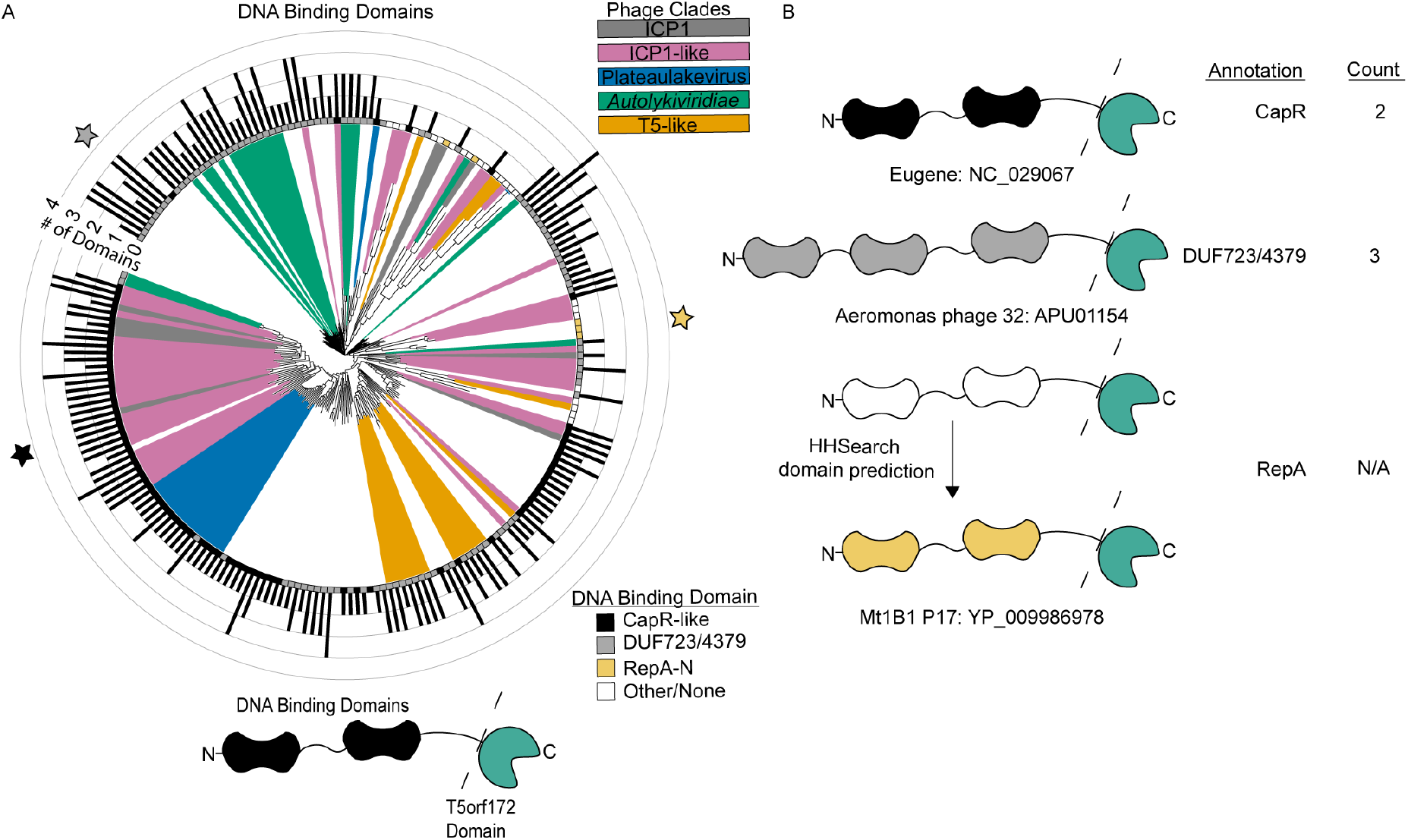
T5orf172 HEGs regularly contain repeated DNA Binding domains from related Zn-finger domains. (A) Alignment of the amino acid sequence of the DNA binding domain(s) of each T5orf172 HEG shows the relationship between the domains. After truncating the amino acid sequence of each T5orf172 HEG to remove the T5orf172 domain, as represented by the dashed line in the model below the figure, the remaining sequence was searched for CapR DNA binding and highly related DUF723 or DUF4397 domains with hmmsearch and boxes at leaf tips were labeled accordingly. The number of DNA binding domains identified by hmmsearch are indicated by the bar graph surrounding the tree. If none of the three Zn-finger domains were identified, the sequence was searched with HHsearch to detect DNA binding domains. Domains identified by HHsearch do not have the number of domains quantified, resulting in a filled-in outer square without a domain occurrence count. Colored stars indicate the representative examples shown in (B), colored according to DNA binding domain type. (B) Representative examples of the domain shading and domain count described in (A). Three representative proteins with CapR, DUF723/4379, or RepA_N annotations in (A). Phage genome names and NCBI protein accession numbers are indicated next to each example.

The modularity of DNA binding regions would lend itself to the rapid diversification of target recognition if modules can be readily exchanged or altered. Beyond their modularity, genes with repeats present on the nucleotide level may provide a favorable substrate for recombination. Even if the level of shared nucleotide identity between repeats is relatively low, recombinants would be expected to arise during phage infection. Generally, characterized phage recombination systems can act on far lower levels of homology and sequence similarity than is required for systems encoded by cellular organisms (61, 62). Using a combination of BLASTn and manual curation, we identified degenerate repeats in the DNA binding coding regions of some ICP1 T5orf172 HEGs (Figure 6A). Being heavily sampled and sequenced, ICP1 isolates provide a clear example of recombination between repeats internal to a HEG, *gp174* (Figure 6A). Recombination between degenerate repeats in ICP1 *gp174* is predicted to have resulted in an in-frame 228bp deletion (Figure 1A, Figure 6A). Previous work has demonstrated that small mutations can cause HEGs to have large shifts in DNA substrate specificity (63). As such, this deletion would be predicted to have a strong effect on HEG specificity.

**Figure 6.**
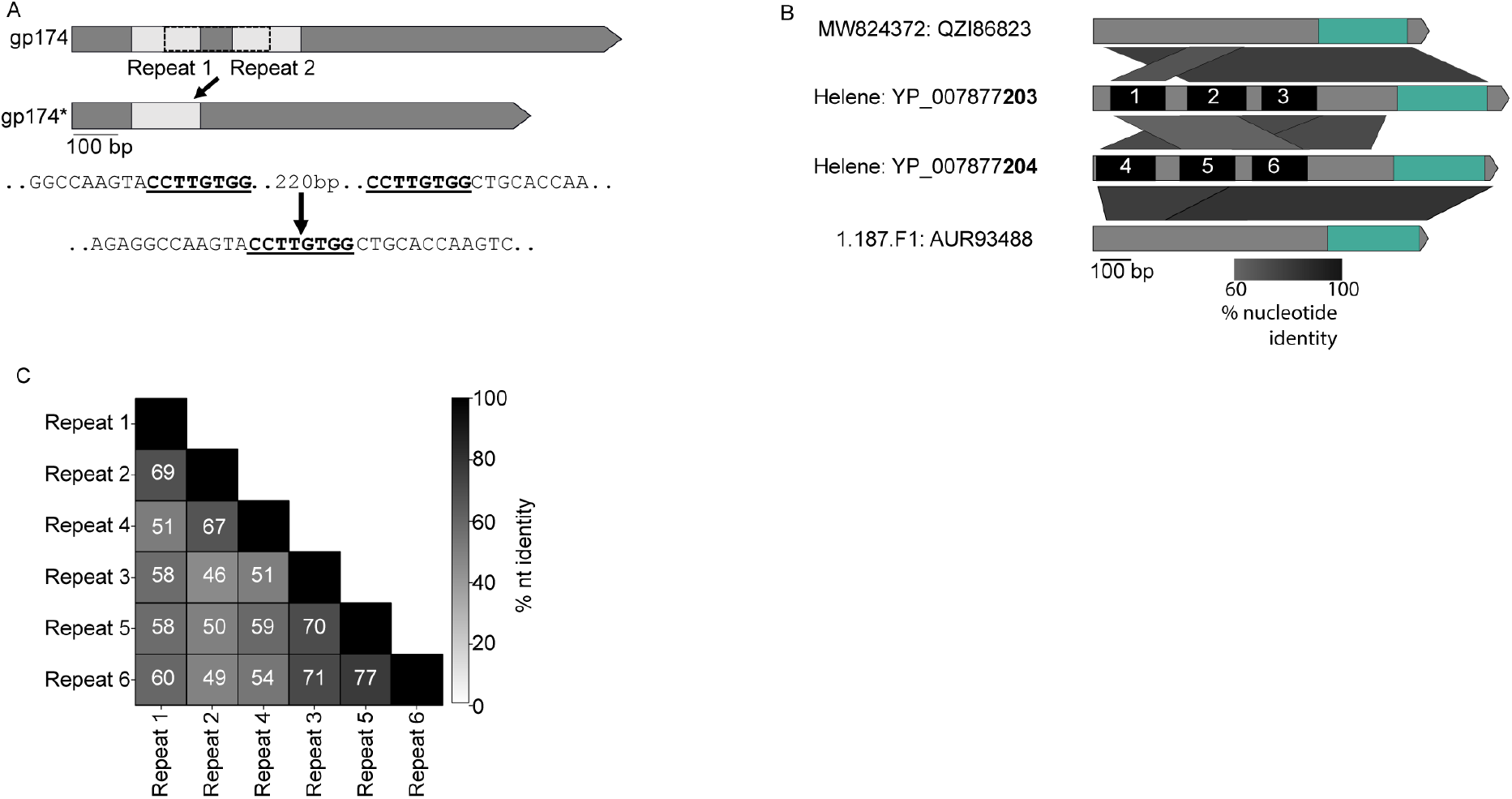
Recombination between HEG DNA binding domains generates sequence duplication. (A) Schematic of predicted recombination events between related repeats in ICP1 *gp174*, a T5orf172 HEG encoding a CapR DNA binding domain. Repeat sequences where predicted recombination occurred are bolded and underlined in the nucleotide sequence below. *gp174\** represents alleles of *gp174* with an in-frame deletion of the repeated sequences. (B) Gene graph comparison of the T5orf172 HEGs from phages Helene, 1.187.F1, and MW824372. The green highlighted region represents the T5orf172 domain of each protein and the black boxes labeled 1-6 represent the repeat domains present in the HEGs in phage Helene. Percent nucleotide identity between the sequences is shown by the shaded parallelograms below the gene graph. (C) A pairwise percent nucleotide identity matrix between the 6 repeat regions of Helene T5orf172 HEGs shown in (B).

In addition to recombination between domains within a single HEG, we also detected evidence for recombination between separate HEG sequences. By generating nucleotide alignments of four HEGs from the Vibrio phages Helene, 1.187.F1, and 184E37.3a (See Table S2 for genome accession numbers), we found evidence highly suggestive of domain swapping between the HEGs. Of these HEGs, a subset contained homologous DNA binding regions but unrelated T5orf172 domains, while others contained unrelated DNA binding regions and homologous T5orf172 domains (Figure 6B and 6C). The homology-driven recombination between HEGs supports that the domains of T5orf172-family HEGs are modular and can be exchanged. A predisposition for recombination is likely to make HEGs more evolvable and may contribute to their success in certain phage lineages.

### HEG-associated domains are engaged in phage-satellite conflicts

Horizontal transfer of HEG-associated domains is not limited to recombination within and between phages. Previous work has also established an evolutionary link between ICP1’s T5orf172 HEGs and genes encoded by PLE, a satellite virus of ICP1. PLE-encoded CapR has similarity to some ICP1-encoded T5orf172 genes and is involved in PLE’s parasitism and antagonism of ICP1 (55). The level of similarity between CapR and some ICP1-encoded T5orf172 HEGs is variable across different PLE variants and appears strongest in a more recently discovered PLE variant, PLE 7. In multiple *capR* alleles, the level of sequence homology to *gp174* is greater than 70% across a 300 bp region, indicating a relatively recent genetic exchange between PLE and ICP1. Notably, both regions of the repeat sequence of *gp174*, as well as another ICP1 T5orf172 HEG *gp165*, share considerable nucleotide identity with *capR* (Figure 7A, 7B), suggesting that similar recombination dynamics could be at play in the evolution of CapR and ICP1 HEGs.

**Figure 7.**
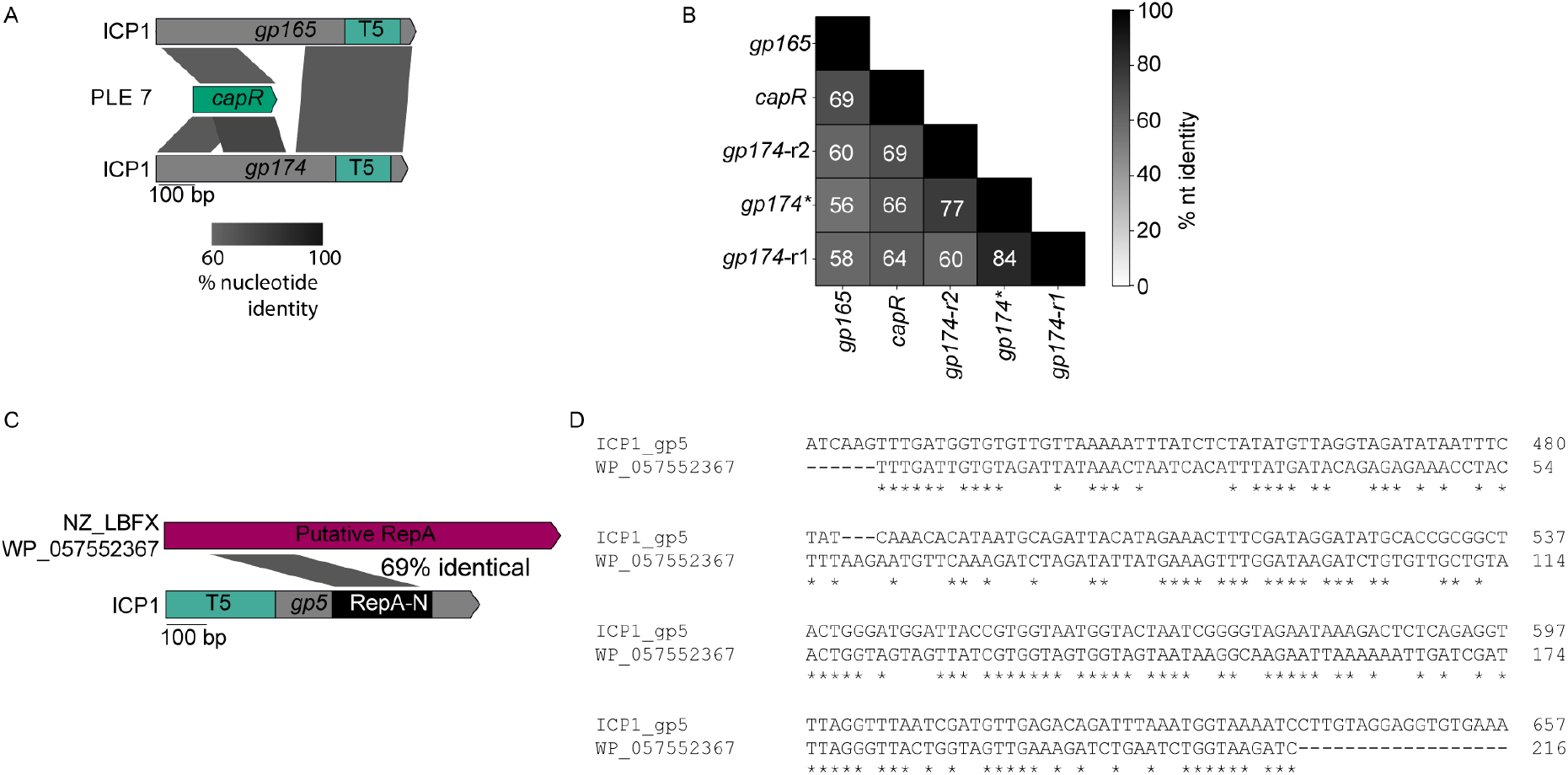
Recombination occurs between satellites and phages. (A) Examples of recombination between ICP1 T5orf172 HEGs and PLE 7 *capR*. Percent nucleotide identity between the sequences is shown by the shaded parallelograms below the gene graph. T5 = T5orf172 domain. (B) A pairwise percent nucleotide identity matrix between the ICP1 T5orf172 HEG *capR* encoding domain and the PLE7 *capR* domain. The two repeat *capR* encoding domains observed in primary *gp174* allele are represented by *gp174-r1* and *gp174-r2*. The ICP1 *gp174\** allele contains in-frame deletion, resulting in a single *capR*-like domain. (C) Recombination between ICP1 T5orf172 HEG *gp5* and non-O1 *V. cholerae* PLE-encoded *repA*. Percent nucleotide identity is shaded between the gp5 RepA-N domain and the putative non-O1 PLE *repA* gene. The protein accession number for the RepA protein is listed to the left of the sequence. (D) A nucleotide alignment revealing the homology highlighted in the diagram in (C). Asterisks indicate identical basepairs between the sequences.

Another characterized example of a HEG-related domain and a horizontally acquired domain being involved in the PLE-ICP1 arms race occurs with ICP1’s <u>o</u>rigin <u>d</u>irected <u>n</u>uclease (Odn). Odn possesses a HEG-derived T5orf172 nuclease domain, and a RepA_N DNA binding domain similar to that of the RepA replication initiation factor encoded by several PLEs (10, 23). During infection, Odn binds and cleaves the cognate PLE origin of replication, preventing PLE from parasitizing ICP1 (10). It is hypothesized that this specificity is an example of HEG domestication, wherein the ancestral Odn allele has recombined with the PLE RepA_N DNA binding domain, increasing the fitness of phages containing Odn as they can inactivate PLE, bolstering phage replication (Figure 8A). Consistent with this, some PLEs possess different RepA_N containing replication initiation factors and cognate origins of replication, and these PLEs are immune to interference by Odn (10). When analyzing the T5orf172 DNA binding domains that lacked one of the three Zn-finger DNA binding domains (CapR, DUF723, or DUF4397) (Figure 5), we found that *odn* was not the only ICP1 gene in the set that encoded a RepA_N domain. A second ICP1 T5orf172 encoding gene, *gp5*, also encodes a RepA_N domain. The RepA_N domain of Gp5 has sequence similarity to the RepA_N domain of a bioinformatically identified MGE with considerable gene synteny to PLE identified from a non-O1 *V. cholerae* isolate (Figure 7C, 7D). The level of amino acid similarity between the Gp5 RepA_N domain and its cognate PLE domain is approximate to the level of similarity between Odn’s RepA_N domain and its own cognate PLE domain. Gp5 is frameshifted in over half of ICP1 isolates (Figure 1A), which we hypothesize indicates that this protein encodes an accessory function for many ICP1 isolates, such as specific defense against PLEs containing this novel RepA/origin combination. These factors suggest that Gp5 may represent a second example of the domestication of a HEG in the ICP1-MGE conflict, resulting in an ICP1 nuclease that is potentially directed against a PLE origin of replication.

**Figure 8.**
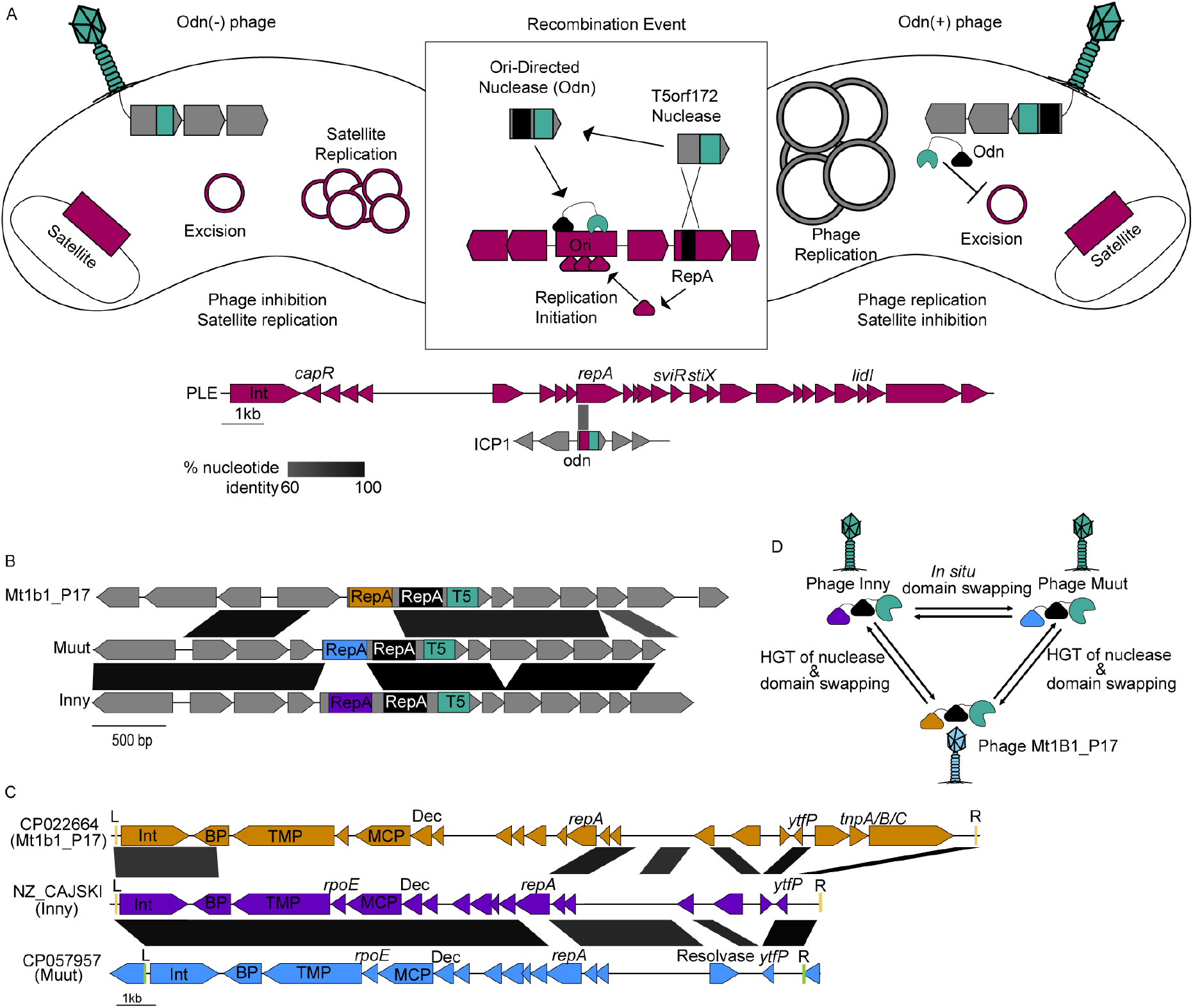
HEGs are broadly involved in the phage-satellite evolutionary arms race. (A) A representative model of the mechanism of satellite inhibition by Odn. In the absence of Odn, the phage satellite life cycle continues unperturbed, inhibiting phage progeny production. Following recombination to acquire Odn, phages are able to overcome the satellite and viable phages are generated from the infection. The PLE gene graph shows the features of PLE 1 and highlights characterized PLE features with labels above the diagram. (B) Examples of phages that are predicted to encode novel *odn* genes. RepA_N domains of putative *odn* genes are colored identically to their corresponding satellite in panel C. Regions of nucleotide identity greater than 60% are shaded accordingly. Each *odn* homolog encodes a second RepA_N domain without detectable homology to cognate satellite *repA* genes, which is shown in black. (C) Putative phage satellites that are predicted to be targets of each respective *odn* homolog indicated in panel B. Gene graphs are colored according to phage genome, which corresponds to the *odn* RepA domain in panel B. Predicted *attL* and *attR* recombinase attachment sites are indicated with red (tRNA integrated) and green (*lysR* integrated) lines and marked L and R. Genes with a strong functional prediction from HHsearch analysis are labeled with gene names or predicted protein function. Dec = decoration, MCP = major capsid protein, Int = integrase, BP = base-plate hub, TMP = tape measure protein. (D) A schematic of the hypothetical exchanges of domains between related phage *odn* homologs.

Aside from the ICP1 RepA_N domain containing T5orf172 proteins, our analysis of DNA binding domains associated with T5orf172 domains (Figure 5) found proteins with similar domain architecture in three closely related *E. coli* phages, Inny, Muut, and Mt1B1_P17 (Figure 8B). These proteins shared a similar domain architecture to Odn, raising the question of whether they may also function as anti-satellite nucleases. Consistent with this hypothesis, a BLASTn search of the DNA encoding the RepA_N domain from each of the three phages against the NCBI nr database detected bacterial homologs of the RepA_N encoding sequence. These bacterial homologs lacked a T5orf172 domain, supporting that they may serve as target sequences for the coliphage encoded T5orf172/RepA_N domain proteins, similar to Odn and PLE replication initiator, RepA (Figure 8A, (10)).

Further inspection of the genomic loci encoding the RepA_N domain homologs revealed characteristic features of phage satellites. These putative satellites encode a similar suite of predicted proteins, are flanked by attachment sites, and lack the full suite of genes for autonomous phage progeny production, suggestive of satellite function (Figure 8C). Similar to the recently described cfPICIs (64), these satellites appear to encode genes required for capsid morphogenesis, such as major capsid proteins and decoration proteins. Nevertheless, these satellites lacked many of the other signature genes of the cfPICI group such as terminase genes, a prohead serine protease, an *alpA* homolog, head-tail adapter genes, and a Pri-Rep replication initiator (64). The new satellites also encode putative tape measure proteins and a predicted component of the phage-tail baseplate but lack the majority of genes required for tail morphogenesis.

Through BLASTn analysis searching the identified putative satellites that match Inny, Muut, and Mt1b1_P17, we were able to identify a set of 224 putative satellites (Table S6) encoded by *E. coli* and closely related organisms. We estimated the ends of most of the putative MGEs by detecting repetitive attachment sites at the end of the elements. Most of the identified elements encode a tyrosine recombinase integrase next to one of the attachment sites and were found integrated next to tRNAs. A subset of the elements, represented by CP057957 (Figure 8C), lacked the most common putative attachment sites and were found to encode additional integration machinery, including a novel serine recombinase and attachment sites. These satellites were found to be occasionally integrated into novel loci compared to the tRNA-proximal elements, suggestive of integration into the novel site by the alternative integration machinery (Table S6).

Much like PLEs, these satellites also exhibited modularity in their RepA N-terminal DNA-binding domain, suggesting the replacement of the satellite replicon components with a functionally analogous nucleotide sequence. Remarkably, this variability in satellite-encoded RepA_N domains was mirrored by the phage nucleases; the homologous T5orf172 domaincontaining proteins in Inny, Muut, and Mt1B1_P17 each encoded separate RepA_N domains, each homologous to a unique satellite RepA protein (Figure 8C, Figure S3A-C). These observations strongly suggest that the three *E. coli* phage nucleases are another outcome of an evolutionary arms race between phages and their satellites and that *odn* genes have evolved multiple times in separate phage lineages, highlighting the neofunctionalization of HEG-associated nuclease domains to the benefit of their genomic hosts.

While the order in which these homologous *odn* genes arose cannot be determined, one can infer reasonable evolutionary steps between them. Inny and Muut have a high level of sequence similarity, while Mt1B1_P17 is much more diverged (Figure S3D). This leads to a model where the change in RepA_N domains between Inny and Muut is an example of domain swapping within a genome, while the nuclease domain must have been horizontally transferred between the Mt1B1_P17 lineage and the Inny and Muut lineage before or after undergoing another DNA binding domain swap (Figure 8D). This model illustrates the flexibility of HEG-associated domains. Capable of both *in situ* mutability and horizontal transfer, these nucleases readily evolve new specificities and functions.

## DISCUSSION

Here we have provided one of the most extensive bioinformatic surveys of phage HEGs to date, with a particular emphasis on a new HEG lineage employing a T5orf172 nuclease domain. Our observations show HEGs are at the vanguard of genetic exchanges, not only between phages but between phage and satellite pairs as well. Of particular note, we find that HEGs bracket essential or otherwise highly selected for gene clusters that are well-conserved at the nucleotide level despite occurring in phage genomes that are related only distantly. We propose these clusters to be a new class of mobile element, the HEG-island. HEG-islands appear to be complex elements composed of two or more flanking HEG genes that provide homing activity and cargo genes that bear key functions of the phage lifecycle. Given the extremely important roles that their cargo genes play, we expect HEG-islands to be a driving force in phage evolution.

Unlike most MGEs, HEG-islands are not expected to have discrete boundaries. HEG-islands are likely to mobilize the totality of their cargo through successive homing events rather than as a single reaction, and partial HEG-island mobilization may explain why some HEG-islands show partial and polar homology to each other (Figure 4). Collaborative homing between flanking HEG sequences would be expected to provide multiple benefits. In a simple scenario, a second proximal HEG could provide a second line of homing activity if one HEG’s target site is absent, or if a competing HEG is homing into the HEG-island. The organization of gene modules into mobile islands that are bracketed by HEGs may be linked to the functional requirements of those modules. The HEG-islands we’ve identified encode for essential heteromeric protein assemblies. Gene conversion of the entire module may be selected for, as conversion of only some genes within a module could bring together alleles that are not functionally compatible, increasing the risk of inviable progeny when only one of the flanking HEGs is able to home into the target sequence. Additionally, multiple HEGs may be a requirement for HEG-island homing between distantly related phages. Homing relies on homologous recombination, and for homing events between distantly related phages, the bulk of the template suitable for homologous recombination would likely be internal to the HEG-islands themselves. In T4, homing is known to be a highly asymmetric process (65). Successive asymmetric recombination events might allow HEG-islands to mobilize into new phage genomes through homology internal to the islands themselves.

HEG-islands complicate the traditional view of HEGs as selfish MGEs. While HEGs are clearly antagonistic towards the alleles they target, homing is mutually beneficial for HEGs and their cargo genes. Organization of cargo into HEG-islands that persist across distantly related phages suggests a high level of coevolution between HEGs and cargo genes. Rather than simply being fortunate bystanders mobilized by the action of a sub-genomic parasite, cargo genes are likely an essential part of HEG-islands, providing homology needed for homing as well as critical functions that select for maintenance of the HEG-island. The turnover of HEG-islands is likely rare and punctuated. Across nearly two decades of sampled ICP1 isolates, we see fairly limited HEG turnover and variation (Figure 1A).

However, analysis of some ICP1-like phage genomes suggests relatively recent exchanges of multiple HEG islands (Figure 4). The exchange of multiple islands encoding essential functions suggests that, at one extreme, some phages can be seen as loose consortiums of HEG-islands which are themselves consortiums of cargo and HEGs.

It is unclear just how ubiquitous HEGs and HEG-islands are among bacteriophages. Our analysis of T5orf172 HEGs and IPA-HNH HEGs reveals several HEG dense phage lineages (Figure 2), while previous work has shown HEGs to be abundant within T4-like phages (6, 14), SPO1-like phages (44), the cluster J phages of *Mycobacterium* (45), and some Staphylococcal phages (43). The work here, while informative, is biased by our initial observations and search parameters, and a more global search for all HEGs encoded within phage genomes would be a rather difficult task. It is hard to predict a phage’s full complement of HEGs. The number of HEG-associated domains has grown with time (53), and in at least one case, a HEG has been found to use a nuclease domain not yet associated with other homing genes (66). Additionally, the most obvious HEG signature, an intronic location, is now known to occur only for a minority of phage HEGs. These factors suggest that this work and prior studies underestimate the abundance of HEGs within phage genomes. Nevertheless, within HEG-dense lineages, some phages clearly possess fewer HEGs, such as the T4-like phage T2, and the ICP1-like phage 1.101.C6 (Figure 1A). The discrepancy in HEG number between related phages suggests that there are fitness costs to encoding HEGs. In the absence of coinfection with partially diverged phages, some HEGs may be selected against to limit the metabolic cost of accessory gene expression. Core genes that evolve without the benefits of homing may also be under stronger selective pressure to perform well in their core functions. Anti-phage defenses that target or detect core phage components (67) could also provide a strong selection for certain core gene alleles, promoting the success of some HEG-less variants. The factors underlying the variable HEG density across bacteriophages are widely unknown.

The relationship between HEGs and their cognate targets and cargo would likely give rise to an arms race dynamic between HEGs, and previous work suggests that HEG arms races do occur (12, 68). In such a context, rapid evolution potentiated by repetitive gene architecture and modular functional domains would be favored. We observe these features among many T5orf172 HEGs, and in at least some cases, we observe recombination between intragenic repeats (Figure 6 and Figure 7). A similar ‘beads on a string’ architecture has been described for the T4 GIY-YIG HEG I-TevI, and is responsible for I-TevI’s functional flexibility as both a homing endonuclease, and a transcriptional repressor. I-TevI is comprised of a C-terminal HTH containing DNA binding domain is linked by a central linker domain to the nucleolytic GIY-YIG domain. This linker domain contains a Zn finger, but rather than functioning as a DNA binding domain, the linker regulates I-TevI’s catalytic activity, allowing I-TevI to function as a transcriptional repressor when no cleavage site is in range of the protein’s binding site (69). Thus, the modular and repetitive architecture of HEG genes likely lends itself towards rapid evolution, functional flexibility, and neofunctionalization through recombination.

HEG gene architecture may also promote the co-option of HEG-related domains for use in phage-satellite conflicts, as has previously been described in ICP1 (10, 55), and as we suggest occurs for a family of *E. coli* phages and their novel satellites (Figure 8). The evolution of the PLE-encoded ICP1 capsid repressor, CapR, from the DNA binding domains of T5orf172 HEGs (55) parallels the evolution of plant AP2 transcription factors from HEGs in bacteria and phages (42). AP2 domains are known to favor GC-rich sequences for targeting (42), and similar biases for GC content or substrate topology could explain why satellites would co-opt HEG DNA binding domains from their helper phage to manipulate that helper phage’s own gene expression. On the side of phage antagonism towards satellites, we see repeated acquisitions of satellite replication initiation factors that are paired with HEG nuclease domains that are expected to facilitate satellite restriction (Figure 8). This likely speaks to satellite origins of replication being effective targets for restriction. The repeated evolution of origin directed nucleases is a notable example of convergent evolution, and highlights the functional versatility of HEG-related genes.

HEG domestication and neofunctionalization pose a limitation to this study. From bioinformatic data alone, it is often not possible to infer whether an individual nuclease gene actively homes. For example, the model coliphage HK97 has been shown to encode an HNH nuclease with domain architecture similar to HNH-HEGs that is directly upstream of the phage terminase and *cos* site. However, this HNH nuclease acts cooperatively with the terminase for cos site cleavage *in vitro* (70). The presence of related HNH nucleases encoded upstream of the terminase in related phages led the authors to conclude that these HNH nucleases are a conserved feature of the phages’ packaging module, while our analysis suggests that the region upstream of their terminase is a hot spot for different classes of HEG-like endonucleases. These two interpretations are not mutually exclusive. Prior work has found that some HEGs bind to and may cut their own sequences at a low rate, possibly to produce a donor DNA end for recombination (12). It is conceivable that this processing could stimulate packaging machinery by altering the topology of the phage genome’s end. Phage termini sequences are often essential for phage propagation, and thus are likely targets for HEG acquisition. The capacity of some HEGs to target their own self-sequence might then allow for the domestication of HEGs as genome end-processing machinery and allow for a gradient of selfish and non-selfish functions among related nuclease genes.

Overall, this work advances our knowledge of phage HEGs while providing a conceptual framework for HEG and HEG-island evolution. We hope the hypotheses presented here can guide future investigations into HEG function and phage evolution. HEGs are at the forefront of horizontal gene transfer between phages, and understanding homing is crucial to understanding how phages evolve.

## DATA AVAILABILITY

Sequence data for samples used for Magic-BLAST analysis can be found in the Sequence Read Archive under the BioProject accession PRJNA609114.

## ACKNOWLEDGEMENT

We would like to acknowledge the members of the Seed Lab for their input during the design and analysis of this manuscript. We would like to thank Zoe Netter for important feedback during the design of the analysis and for helping with revising the manuscript.

