## Supplementary material for "Nuclease genes occupy boundaries of genetic exchange between bacteriophages": Figures S1-S3

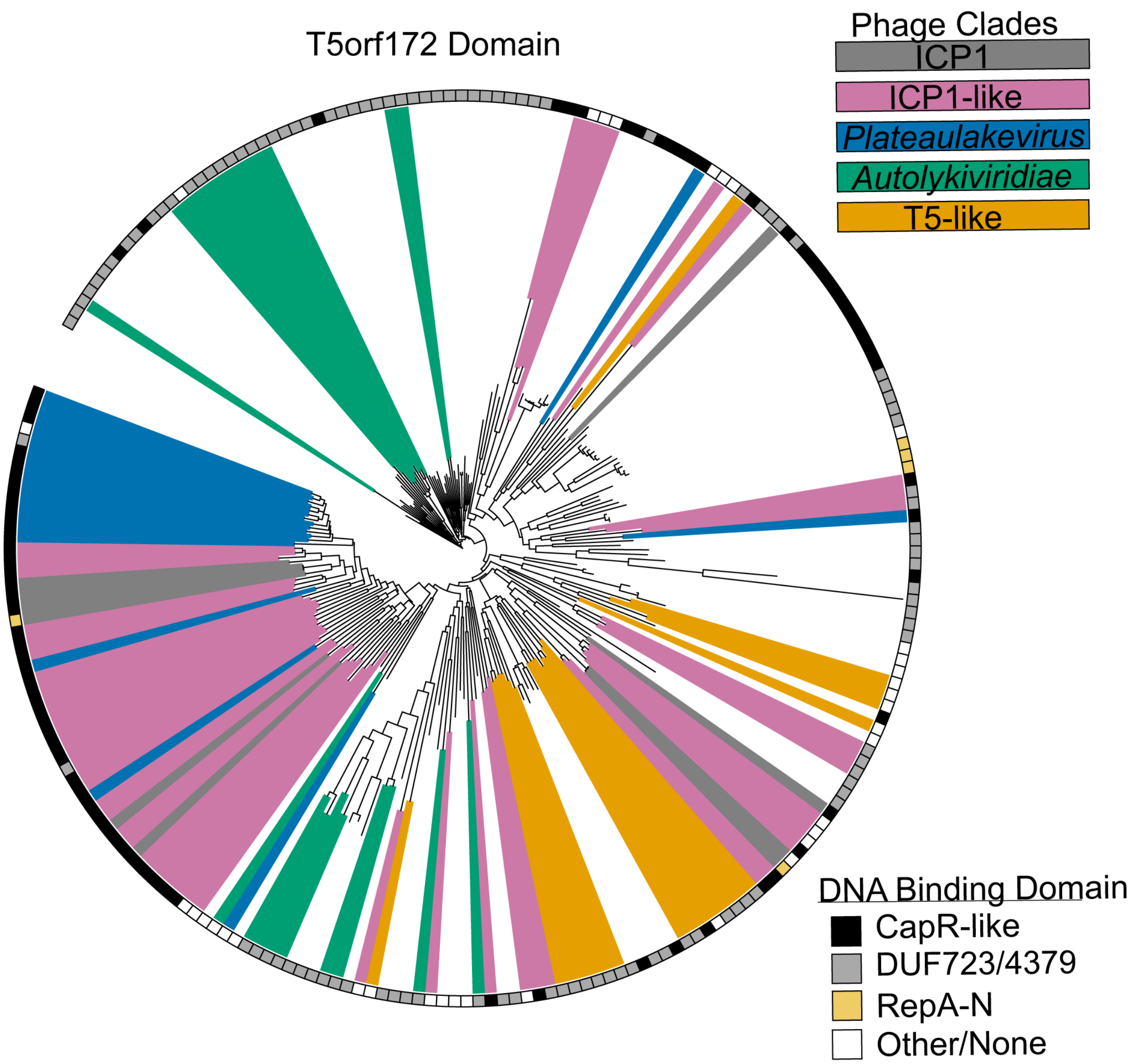


**Figure S1. Phylogenetic analysis of the distinct T5orf172 domains of phages**
Alignment of the T5orf172 domains from each of the identified T5orf172 HEG-containing phages. Phage clades are determined based on the phylogenetic tree in Figure 2. The ring of boxes surrounding the tree corresponds to the identified DNA binding domain for each individually graphed HEG, as shown in Figure 5.

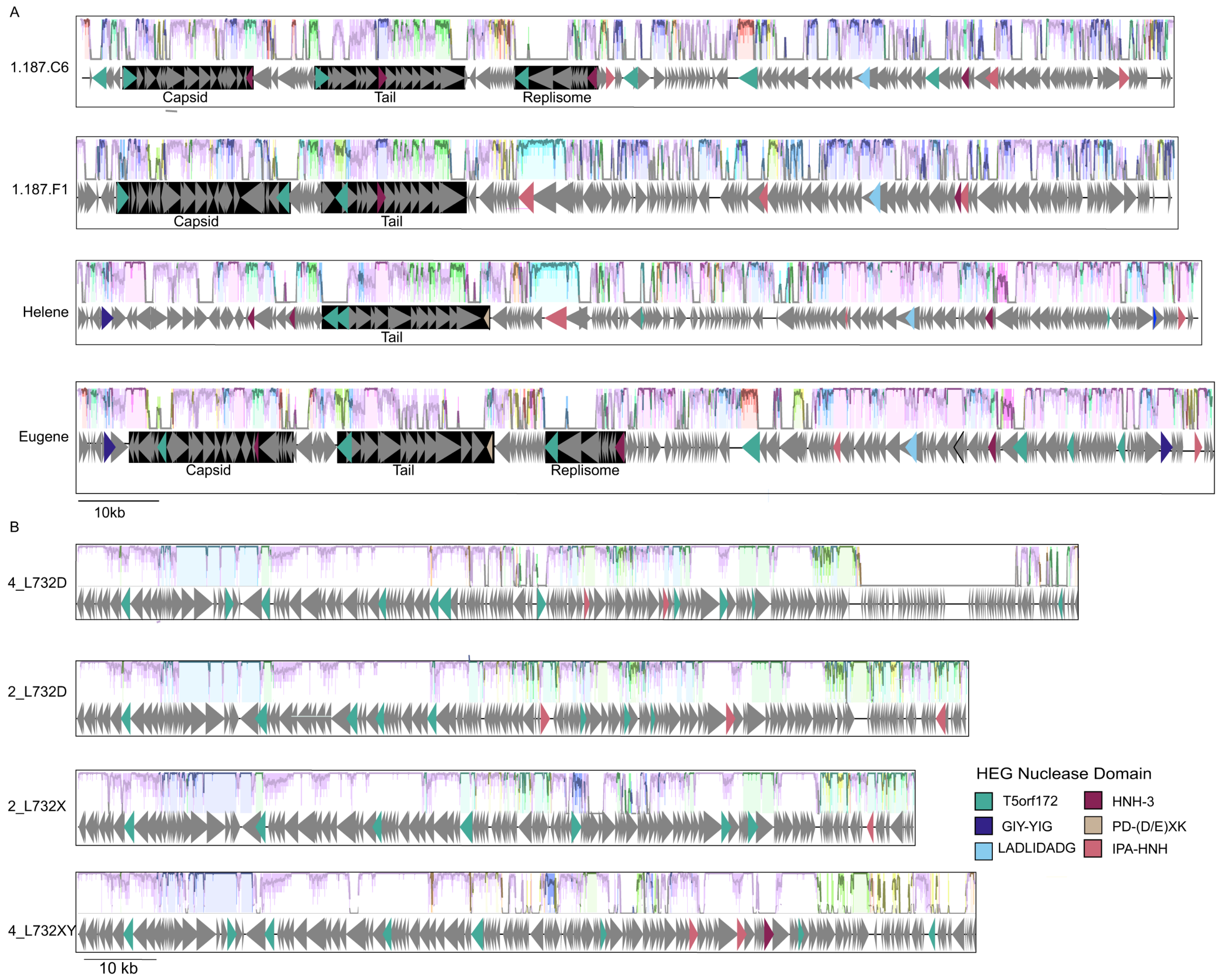

A
1.187.C6
Capsid
Tail
Replisome
1.187.F1
Capsid
Tail
Helene
Tail
Eugene
Capsid
Tail
Replisome
10kb
B
4_L732D
2_L732D
2_L732X
4_L732XY
10 kb
HEG Nuclease Domain
T5orf172
HNH-3
GIY-YIG
PD-(D/E)XK
LADLIDADG
IPA-HNH

**Figure S2. Related HEG-Islands containing cargo genes are flanked by variable HEG families in related phages**

Whole-genome alignment of ICP1-like (A) and *Plateaulikevirus* (B) generated with Mauve (https://journals.plos.org/plosone/article?id=10.1371/journal.pone.0011147 ) shows the relatedness of HEG-islands in otherwise distinct phages. Genes encoding predicted HEG nuclease domains are colored according to the key, and gene neighborhoods highlighted in Figure 4 are shaded in black. Despite sharing conserved cargo genes, many gene neighborhoods are flanked by HEGs with different nuclease domains.

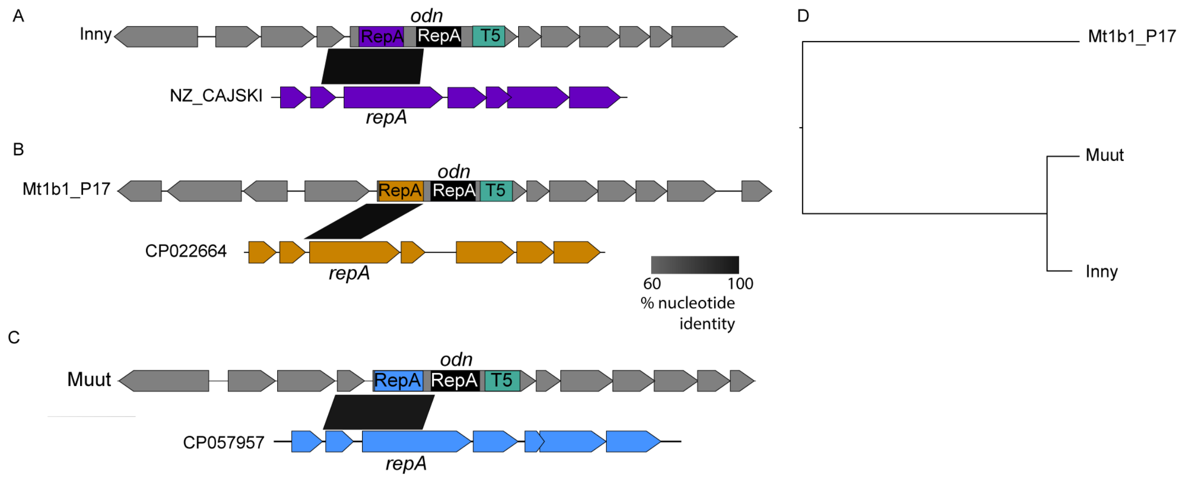


**Figure S3. Recombination of HEG-associated domains between phages and their satellites.**

(A-C) Examples of recombination between *E. coli* phages shown in Figure 8 and their satellites, specifically highlighting the RepA-N domain homology of the satellites to the respective phages. Regions of > 60% identity are marked according to the legend with black parallelograms.

(D) VipTreeGen phylogeny of each of the three phage genomes, showing the variable conservation of coding sequences between the phages
