## Supplementary material for "Nuclease genes occupy boundaries of genetic exchange between bacteriophages": Tables S1-S3

Table S1) *Vibrio cholerae* and ICP1 Isolates used in this study
Accession information for genomes of *Vibrio cholerae* and ICP1 references used throughout this study

| Strain | Accession | Genome Citation |
|---|---|---|
| *Vibrio cholerae* E7946 | CP024162, CP024163 | Camilli, A. 2017 |
| ICP1 1992_Ind_M4 | MW794141 | Boyd et al., 2021 |
| ICP1 2001_Dha_0 | HQ641347 | Seed et al., 2011 |
| ICP1 2001_Dha_A | HQ641353 | Seed et al., 2011 |
| ICP1 2003_Dha_A | MW794140 | Boyd et al., 2021 |
| ICP1 2004_Dha_A | HQ641354 | Seed et al., 2011 |
| ICP1 2005_Dha_A | HQ641352 | Seed et al., 2011 |
| ICP1 2006_Dha_A | HQ641351 | Seed et al., 2011 |
| ICP1 2006_Dha_B | HQ641350 | Seed et al., 2011 |
| ICP1 2006_Dha_C | HQ641349 | Seed et al., 2011 |
| ICP1 2006_Dha_D | HQ641348 | Seed et al., 2011 |
| ICP1 2006_Dha_E | MH310934 | Angermeyer et al., 2018 |
| ICP1 2006_Dha_EΔCR_ ΔCas2_3 | N/A | McKitterick and Seed, 2018 |
| ICP1 2011_Dha_A | MH310933 | Angermeyer et al., 2018 |
| ICP1 2011_Dha_B | MH310935 | Angermeyer et al., 2018 |
| ICP1 2012_Ind_A | MH310936 | Angermeyer et al., 2018 |
| ICP1 2015_Dha_A | MW794150 | Boyd et al., 2021 |
| ICP1 2016_Dha_A | MW794151 | LeGault et al., 2021 |
| ICP1 2017_Dha_A | MW794152 | LeGault et al., 2021 |
| ICP1 2017_Dha_AA | MW794153 | LeGault et al., 2021 |
| ICP1 2017_Dha_AB | MW794154 | LeGault et al., 2021 |
| ICP1 2017_Dha_AC | MW794155 | LeGault et al., 2021 |
| ICP1 2017_Dha_AD | MW794156 | LeGault et al., 2021 |
| ICP1 2017_Dha_AE | MW794157 | LeGault et al., 2021 |
| ICP1 2017_Dha_B | MW794158 | LeGault et al., 2021 |
| ICP1 2017_Dha_C | MW794159 | LeGault et al., 2021 |
| ICP1 2017_Dha_D | MW794160 | LeGault et al., 2021 |
| ICP1 2017_Dha_E | MW794161 | LeGault et al., 2021 |
| ICP1 2017_Dha_F | MN419153 | LeGault et al., 2021 |
| ICP1 2017_Dha_N | MW794162 | LeGault et al., 2021 |
| ICP1 2017_Dha_O | MW794163 | LeGault et al., 2021 |
| ICP1 2017_Dha_P | MW794164 | LeGault et al., 2021 |
| ICP1 2017_Dha_R | MW794165 | LeGault et al., 2021 |
| ICP1 2017_Dha_S | MW794166 | LeGault et al., 2021 |
| ICP1 2017_Dha_V | MW794167 | LeGault et al., 2021 |

| | | |
|---|---|---|
| ICP1 2017_Dha_W | MW794168 | LeGault et al., 2021 |
| ICP1 2017_Dha_X | MW794169 | LeGault et al., 2021 |
| ICP1 2017_Dha_Y | MW794170 | LeGault et al., 2021 |
| ICP1 2017_Dha_Z | MW794171 | LeGault et al., 2021 |
| ICP1 2017_DRC_106 | MW794142 | Alam et al., 2022 |
| ICP1 2017_DRC_32 | MW794143 | Alam et al., 2022 |
| ICP1 2017_DRC_48 | MW794144 | Alam et al., 2022 |
| ICP1 2017_DRC_55 | MW794145 | Alam et al., 2022 |
| ICP1 2017_DRC_72 | MW794146 | Alam et al., 2022 |
| ICP1 2017_DRC_74 | MW794147 | Alam et al., 2022 |
| ICP1 2017_DRC_82 | MW794148 | Alam et al., 2022 |
| ICP1 2017_DRC_87 | MW794149 | Alam et al., 2022 |
| ICP1 2017_Mat_H | MN419153 | LeGault et al., 2021 |
| ICP1 2017_Mat_I | MW794172 | LeGault et al., 2021 |
| ICP1 2017_Mat_K | MW794173 | LeGault et al., 2021 |
| ICP1 2018_Mat_001 | MW794174 | LeGault et al., 2021 |
| ICP1 2018_Mat_002 | MW794175 | LeGault et al., 2021 |
| ICP1 2018_Mat_004 | MW794176 | LeGault et al., 2021 |
| ICP1 2018_Mat_159 | MW794177 | LeGault et al., 2021 |
| ICP1 2018_Mat_160 | MW794178 | LeGault et al., 2021 |
| ICP1 2018_Mat_164 | MW794179 | LeGault et al., 2021 |
| ICP1 2018_Mat_166 | MW794180 | LeGault et al., 2021 |
| ICP1 2018_Mat_167 | MW794181 | LeGault et al., 2021 |
| ICP1 2018_Mat_170 | MW794182 | LeGault et al., 2021 |
| ICP1 2018_Mat_B | MW794183 | LeGault et al., 2021 |
| ICP1 2019_Dha_007 | MW794184 | LeGault et al., 2021 |
| ICP1 2019_Dha_G | MW794185 | LeGault et al., 2021 |
| ICP1 2019_Dha_H | MW794186 | LeGault et al., 2021 |
| ICP1 2019_Dha_I | MW794187 | LeGault et al., 2021 |
| ICP1 2019_Mat_005 | MW794188 | LeGault et al., 2021 |
| ICP1 2019_Mat_B | MW794189 | LeGault et al., 2021 |
| ICP1 2019_Mat_C | MW794190 | LeGault et al., 2021 |
| ICP1 2019_Mat_D | MW794191 | LeGault et al., 2021 |
| ICP1 2019_Mat_E | MW794192 | LeGault et al., 2021 |

Table S2) Other organisms referenced in this study
The accession numbers and shorthand notation for all strains referred to in the study.

| Accession | Shortened name[a] | Common name | Host | Genome Citation |
|---|---|---|---|---|
| AP014858 | RYC | Vibrio phage RYC | *Vibrio coralliilyticus* | Ramphul et al. 2017 |
| HQ316579 | Helene | Vibrio phage helene 12B3 | *Vibrio splendidus* | NA[b] |
| HQ634156 | PWH3a-P1 | Vibrio phage PWH3a-P1 | *Vibrio natriegens* | NA |
| HQ634195 | Eugene | Vibrio phage eugene 12A10 | *Vibrio spp.* | NA |
| KX507046 | S4-7 | Vibrio phage S4-7 | *Vibrio anguillarum* | NA |
| NC_047839 | SL20 | Pseudoalteromonas phage SL20 | *Pseudoaltero monas spp.* | NA |
| NC_048769 | 2_L372D | Aeromonas phage 2-L372D | *Aeromonas hydrophila* | NA |
| NC_048770 | 2_L372X | Aeromonas phage 2-L372X | *Aeromonas hydrophila* | NA |
| NC_048771 | 4_L372D | Aeromonas phage 4_L372D | *Aeromonas hydrophila* | NA |
| NC_048772 | 4_L372XY | Aeromonas phage 4_L372XY | *Aeromonas hydrophila* | NA |
| MG592529 | 1.161.C5 | Vibrio phage 1.161.O._10N.261.48.C5 | *Vibrio lentus* | Kauffman et al. 2018 |
| MG592562 | 1.191.C6 | Vibrio phage 1.193.O._10N.286.52.C6 | *Vibrio splendidus* | Kauffman et al. 2018 |
| MG592473 | 1.101.C6 | Vibrio phage 1.101.O._10N.261.45.C6 | *Enterovibrio norvegicus* | Kauffman et al. 2018 |
| MG592553 | 1.187.F1 | Vibrio phage 1.187.O._10N.286.49.F1 | *Vibrio splendidus* | Kauffman et al. 2018 |
| AP018813 | T2 | Enterobacteria phage T2 | *Escherichia coli* | Akiyama et al. 2018 |
| NC_000866 | T4 | Enterobacteria phage T4 | *Escherichia coli* | Miller et al. 2003 |
| NC_020843 | 11895-B1 | Vibrio phage 11895-B1 | *Vibrio spp.* | NA |
| NC_025436 | 1/4 | Shewanella sp. phage 1/4 | *Shewanella spp.* | Senčilo et al. 2015 |
| NC_025470 | 1/40 | Shewanella sp. phage 1/40 | *Shewanella spp.* | Senčilo et al. 2015 |
| NC_029057 | qdvp001 | Vibrio phage qdvp001 | *Vibrio spp.* | NA |
| MK719750 | Barba31A | Rheinheimera phage vB_RspM_Barba31A | *Rheinheimer a spp.* | Nilson et al. 2019 |
| MK719708 | Barba4S | Rheinheimera phage vB_RspM_Barba4S | *Rheinheimer a spp.* | Nilson et al. 2019 |
| NC_030934 | PsyM_Kil1 | Pseudomonas phage vB_PsyM_KIL1 | *Pseudomona s syringae* | Rombouts et al. 2016 |
| NC_052657 | Muut | Escherichia phage muut | *Escherichia coli* | Olsen et al. 2020 |

| | | | | |
|---|---|---|---|---|
| MN850601 | Inny | Escherichia phage inny | *Escherichia coli* | Olsen et al. 2020 |
| NC_052662 | Mt1B1_P17 | Escherichia phage Mt1B1_P17 | *Escherichia coli* | NA |
| CP022664 | CP022664 | Escherichia coli strain FORC 064 chromosome | Bacterial | NA |
| NZ_CAJSKI 010000035 | NZ_CAJSKI | Escherichia coli isolate Fecal samples | Bacterial | NA |
| NZ_LBFX01 000018 | YB2A06 | Vibrio cholerae strain YB2A06 | Bacterial | NA |
| CWPX0100 0013 | PLE7 *V. cholerae* | Vibrio cholerae genome assembly 4056_7#9, scaffold ERS013187SCcontig000013 | Bacterial | NA |
| CP057957 | N/A | Escherichia coli strain RHB08-C21 chromosome | Bacterial | AbuOun et at. 2021 |

[a]Phage names shortened for simplicity in figure and text.

[b]Abbreviation: NA, not applicable.

Table S3. Pfam domains used for gene neighborhood analysis

| **Family** | **Pfam Domain(s)** |
|---|---|
| Capsid | PF03864, PF05065, PF05357, PF07068 |
| Terminase | PF03592,PF03237,PF04466,PF07471,PF11053,PF16677,PF17288,PF17289,PF05876,PF05944,PF06056,PF03354,PF05119 |
| Tape Measure | PF05017, PF06120, PF06791, PF09718, PF10145, PF20155, PF16459, PF16460, PF16461,PF17388,PF19268 |
| Ribonucleotide Reductase | PF14597,PF00268, PF00317,PF02867,PF08343 |
| DNA Polymerase | PF00476, PF02767 |
| T5orf172 | PF10544,PF13455 |
| GIY-YIG | PF01541 |
| HNH-3 | PF01381 |
| LAGLIDADG | PF14528,PF03161,PF00961 |
| CapR | Custom profile |
| IPA-HNH | Custom profile |
